# Early-life adversity alters adult nucleus incertus neurons: implications for neuronal mechanisms of increased stress and compulsive behavior vulnerability

**DOI:** 10.1101/2024.12.02.626443

**Authors:** Anna Gugula, Patryk Sambak, Aleksandra Trenk, Sylwia Drabik, Aleksandra Nogaj, Zbigniew Soltys, Andrew L. Gundlach, Anna Blasiak

## Abstract

**BACKGROUND:** Early-life stress (ELS) arising from physical and emotional abuse disrupts normal brain development and impairs hypothalamic-pituitary-adrenal axis function, increasing the risk of psychopathological disorders and compulsive behaviors in adulthood. However, the underlying neural mechanisms remain unclear. The brainstem nucleus incertus (NI) is a highly stress-sensitive locus, involved in behavioral activation and stress-induced reward (food/alcohol) seeking, but its sensitivity to ELS remains unexplored.

**METHODS:** We used neonatal maternal separation stress in rats as a model for ELS and examined its impact on stress-related mRNA and neuropeptide expression in the NI, using fluorescent *in situ* hybridization and immunohistochemistry, respectively. Using whole-cell, patch-clamp recordings we determined the influence of ELS on the synaptic activity, excitability, and electrophysiological properties of NI neurons. Using c-Fos protein expression we also assessed the impact of ELS on the sensitivity of NI neurons to acute restraint stress in adulthood.

**RESULTS:** ELS weakened the acute stress responsiveness of NI neurons, and caused dendritic shrinkage, impaired synaptic transmission and altered electrophysiological properties of NI neurons in a cell-type-specific manner. Additionally, ELS increased the expression of mRNA encoding corticotropin-releasing hormone receptor type 1 and the nerve-growth factor receptor, TrkA in adult NI.

**CONCLUSIONS:** The multiple, cell-type specific changes in the expression of neuropeptides and molecules associated with stress and substance abuse in the NI, as well as impairments in NI neuron morphology and electrophysiology caused by early-life stress and observed in the adult brain, may contribute to the increased susceptibility to stress and compulsive behaviors observed in individuals with a history of ELS.

## INTRODUCTION

Early-life stress (ELS) arising from physical and emotional abuse, disrupts normal brain development and impairs hypothalamic–pituitary–adrenal (HPA) axis function, and a large body of evidence has shown that children who experience such adversities have an elevated risk of developing psychopathological disorders in adulthood (1–3). Moreover, a common consequence of experiencing ELS is increased susceptibility to compulsive behaviors, such as compulsive food consumption and addiction to psychoactive substances (4–6). Furthermore, ELS increases stress sensitivity in adulthood and exposure to severe stressors throughout life can markedly increase the likelihood of developing both psychopathological disorders and substance abuse in individuals who have experienced ELS (7–12). However, although the consequences of adverse experiences in early life associated with the development of mental disorders and substance abuse have been widely described, the underlying neural mechanisms are still largely unknown.

An area of the brain that is currently unexplored in the context of ELS sensitivity, is the brainstem nucleus incertus (NI), a highly stress-sensitive and conserved component of an ascending arousal network, described in human (13), macaque (14), rat (15), mouse (16) and fish (17) brain. The NI contains heterogeneous populations of mainly GABA, but also glutamate, neurons, that give rise to ascending forebrain projections, and it is the major source of the neuropeptide, relaxin-3 (RLN3) in the brain (15,18–21). Considerable pharmacological and functional data indicate an important role of the NI and RLN3 in stress responses and behavioral activation (15,22–29), as well as compulsive behaviors and stress-induced reward seeking (30–34). Human and animal studies have shown that NI neurons abundantly express stress-related corticotropin-releasing hormone receptor type 1 (CRHR1), and virtually all RLN3-positive NI neurons co-express CRHR1 (27,35,36). RLN3 and its cognate receptor, RXFP3 (relaxin family peptide 3 receptor), control stress-induced alcohol preference in mice (32) and reinstatement of alcohol seeking in rats (30,31,36,37), via CRHR1 signaling in the NI (27,38). Importantly, the potent effects of ELS on the expression of CRH and CRHR1, as well as the tight link between this influence and increased susceptibility to developing substance abuse have been repeatedly demonstrated in humans and animals (39,40), yet the possible influence of ELS on NI neurons, including CRHR1 expression, remains unknown. Notably, in addition to RLN3, NI neurons synthesize other neuropeptides, including cholecystokinin (CCK) (15,41). Our previous studies have demonstrated that CCK is synthesized by a separate neuronal population to that synthesizing RLN3 (15), but the vulnerability of NI CCK neurons to different stressors, including ELS and acute stress remains to be elucidated.

The detrimental consequences of ELS also involve disturbances in nerve-growth factor (NGF)-mediated tropomyosin receptor kinase A (TrkA) signaling (42,43). NGF/TrkA signaling, in addition to its canonical trophic actions, is strongly implicated in stress responses, and exerts stimulatory actions on the HPA axis. Furthermore, NGF levels are sensitive to stressful events (44–46), and NGF/TrkA signaling is strongly implicated in alcohol and other drug use disorders (47,48), but the underlying neuronal mechanisms are not well understood. Notably, NI neurons in the rat abundantly express TrkA (49,50), yet to our knowledge, there is no data regarding the influence of stress on TrkA expression in the brainstem, including the NI.

Therefore, in the current study, we examined the influence of neonatal maternal separation (MS), a well-documented animal model of ELS, on the expression of specific neuropeptide and stress-related receptor mRNA species in GABA and glutamate NI neurons in male rats. We also examined the impact of MS on the synaptic activity, excitability, and membrane electrophysiological properties of defined NI neurons, and the consequences for NI neuronal morphology. Finally, we determined the influence of MS on the sensitivity of NI neurons to acute stress in adulthood.

## MATERIALS AND METHODS

All experiments were approved by the 2^nd^ Local Institutional Animal Care and Use Committee (Krakow, Poland), and conducted according to the EU Directive 2010/63/EU on the protection of animals used for scientific purposes.

Male Sprague-Dawley rat pups from the experimental group were subjected to an MS procedure – a daily 3-h separation from dams during postnatal days (PNDs) 2–14. Adult male rats were used in all experiments. In the first experiment, rats were subjected to an acute restraint stress, and transcardially perfused for immunohistochemistry. Coronal sections throughout the NI were immunostained and c-Fos protein levels in the NI were assessed as a marker of neuronal activation. A second series of experiments involved whole-cell current and voltage patch-clamp recordings, and recorded NI neurons were filled with biocytin, immunostained, and their dendritic morphology was examined. In a third experiment, RNAscope HiPlex *in situ* hybridization was performed.

A detailed description of all procedures is provided in the Supplement.

## RESULTS

### MS and restraint stress differentially affected c-Fos expression in NI neurons

In studies to determine the impact of MS on the sensitivity of NI neurons to acute stress in adulthood, rats (8 weeks old) from control (Ctrl) and MS cohorts were divided into groups that were subjected to restraint stress (S) or not (NS). Activation of relaxin-3 (RLN3+) and pro-cholecystokinin (pCCK+) positive NI neurons was assessed in these groups by the presence of c-Fos protein. RLN3+ and pCCK+ neurons with (+) and without (−) c-Fos were counted (RLN3+/c-Fos+, pCCK+/c-Fos+ and RLN3+/c-Fos–, pCCK+/c-Fos–) (Figure 1A-C). A semi-automated analysis of binarized signal from NI-located c-Fos+ nuclei was also performed.

**Figure 1.**
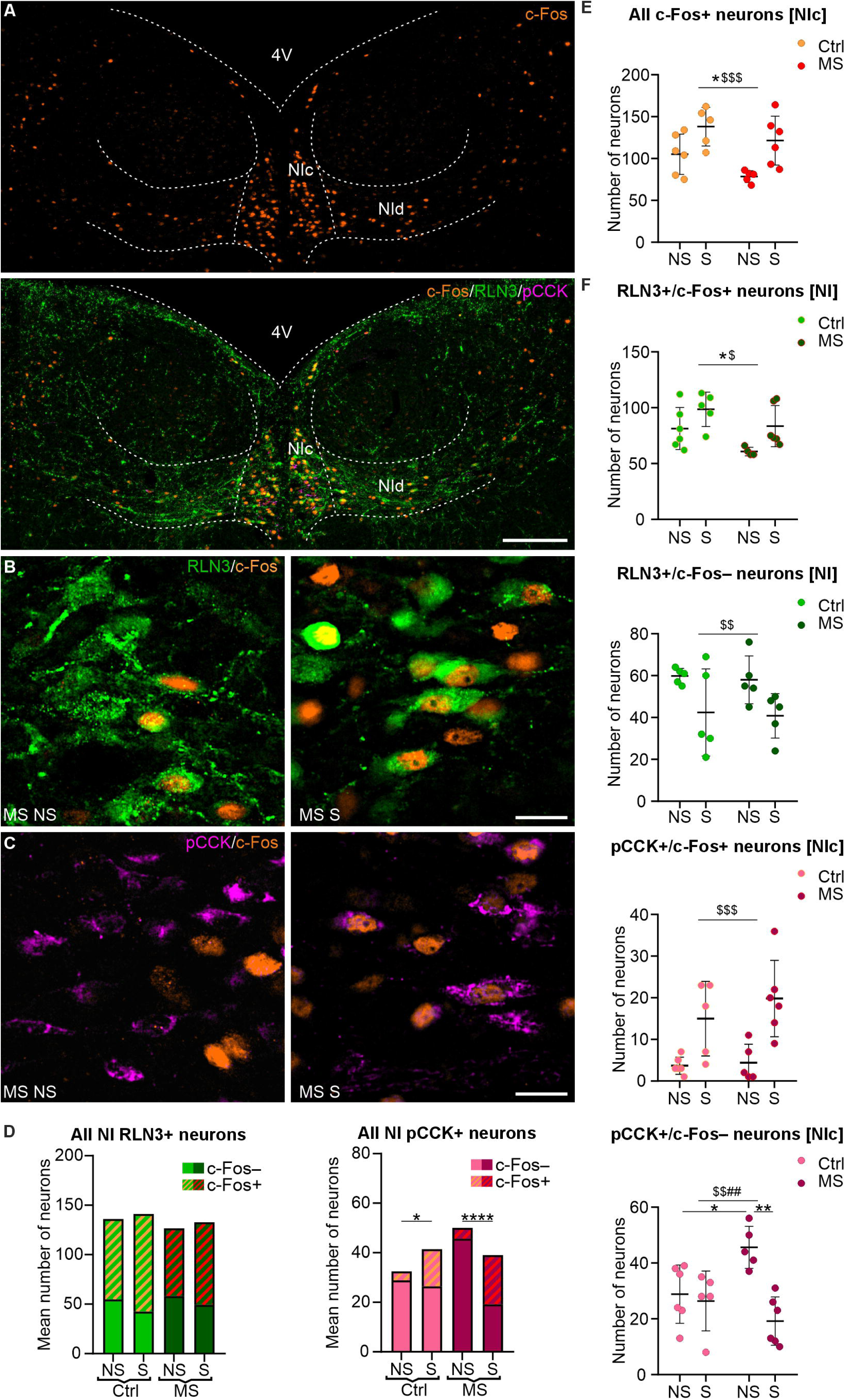
Maternal separation and restraint stress differentially modify the number of c-Fos-expressing NI neurons. **(A)** Representative images of c-Fos protein expression in NI. Scale bar: 200 μm. **(B, C)** Magnified NIc area with RLN3-(**B**) and pCCK-immunopositive (**C**) NI neurons expressing c-Fos in MS NS (left) and MS S (right) rats. Scale bar: 20 μm. (**D)** Proportions of RLN3-positive (RLN3+, left) and pCCK-positive (pCCK+, right) NI c-Fos+ and c-Fos– neurons in all tested groups of rats: control (Ctrl) and MS, subjected (S) and not subjected to restraint stress (NS). Note that both MS and restraint stress altered these proportions in pCCK– but not RLN3+ cells. Statistical significance was determined using Fisher’s exact test: * (p<0.05), **** (p<0.0001). Please see **Supplementary Table 2** for details. **(E)** The number of all NIc c-Fos+ cells was increased by restraint stress and decreased by MS under both conditions (NS and S). **(F)** MS and restraint stress differentially influenced the number of neurochemically defined c-Fos+ and c-Fos– NI/NIc neurons. Statistical significance in **(E)** and **(F)** was determined using two-way ANOVA with a post hoc Tukey test: * MS effect, $ restraint stress effect, # interaction of MS and restraint stress. The number of each symbol indicates the level of statistical significance: * (p<0.05), ** (p<0.01), *** (p<0.001). Please see **Supplementary Table 1** for details. Abbreviations: 4V, 4th ventricle; NI, nucleus incertus; NIc, nucleus incertus pars compacta; NId, nucleus incertus pars dissipata; MS, maternal separation; pCCK, pro-cholecystokinin; RLN3, relaxin-3.

Both MS and restraint stress altered the total number of c-Fos-expressing cells in the NI pars compacta (NIc). Expression of c-Fos in NIc was increased by restraint and decreased by MS in both groups (Ctrl S and MS S; Figure 1E, Supplementary Table 1). An effect of restraint stress was also observed in the NI pars dissipata (NId), and when the whole NI area was analyzed (Supplementary Figure 1).

Cell counting and subsequent ANOVA revealed that restraint stress caused an increase in the number of RLN3+/c-Fos+ cells in the NI in both the Ctrl and MS group, which was accompanied by a decrease in the number of RLN3+/c-Fos– neurons (Figure 1F, Supplementary Table 1).

Importantly, in MS rats significantly fewer RLN3+ neurons expressed c-Fos in both the S and NS groups (Figure 1F, Supplementary Table 1). The effects of restraint stress were significant in NIc, but not NId (Supplementary Figure 1, Supplementary Table 1).

Counting of pCCK+ neurons was performed only in NIc, as CCK in the NI is synthesized almost exclusively by neurons located in this area (15). Similar to the effect on RLN3+ cells, ANOVA indicated an increase in the number of pCCK+/c-Fos+ neurons after restraint stress in both Ctrl and MS groups, relative to the NS groups (Figure 1F, Supplementary Table 1). Notably, a significant effect of restraint stress and an interaction with MS was detected in pCCK+/c-Fos– NIc cells. Post-hoc analysis revealed that MS produced a significant increase in the number of pCCK+/c-Fos– neurons, and restraint stress in the MS group reversed this effect (Figure 1F, Supplementary Table 1).

Notably, restraint stress altered the proportions of c-Fos+ and c-Fos– pCCK+ neurons, towards a higher level of c-Fos expression. The effect was significant in both the Ctrl and MS group, but more pronounced in the latter (Figure 1D, Supplementary Table 2).

### MS altered the electrophysiology of adult NI neurons

#### MS altered the active and passive membrane properties of NI neurons

In studies to investigate the impact of MS on the passive and active membrane properties of NI neurons, a series of *ex vivo* whole-cell patch-clamp experiments was conducted in current clamp mode (holding potential –75 mV). Immunohistochemical procedures combined with anti-biocytin staining, that followed these electrophysiological experiments, allowed the categorization of examined neurons based on their spatial distribution (NIc and NId) and peptide content (RLN3+ NI neurons and pCCK+ NI neurons) (Figure 2A). The electrophysiological type of the recorded neurons was determined based on presence (type I) or absence (type II) of an A-type potassium current, as described (19) (Supplementary Figure 2).

**Figure 2.**
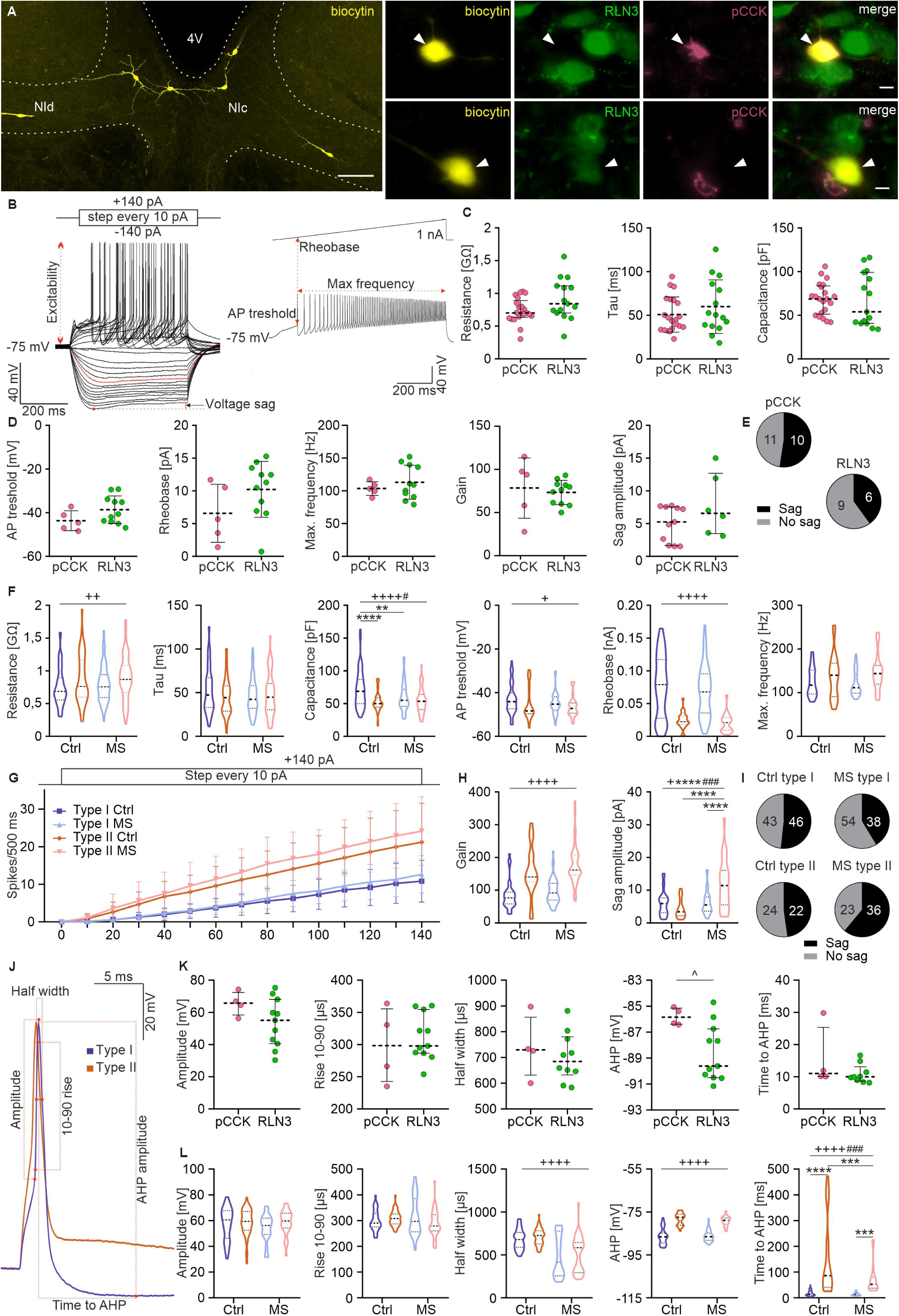
Influence of electrophysiological type and MS on passive and active neural properties of NI neurons. **(A)** Immunohistochemical localization of recorded neurons in the NI, filled with biocytin (left side, scale bar: 100 μm), and fluorescent projection images of biocytin-filled pCCK+ (top) or RLN3+ (bottom) NI neurons, indicated by arrowheads (scale bar: 20 μm). **(B)** Current step stimulation protocol (top left) and voltage responses of an exemplary NI neuron (bottom left), illustrating the measured neural properties. Neuronal excitability (input-output relationship) was assessed by counting the number of APs evoked by depolarizing current pulses (0–140 pA). Membrane resistance, time constant, and capacitance were measured from the voltage response to a –50 pA hyperpolarizing current pulse. Resistance was calculated by subtracting the voltage current from the ohmic current-voltage relation; the time constant was measured in the charging phase of the voltage response, and capacitance was calculated by dividing the resistance and time constant. Voltage sag was measured from the voltage response to a –140 pA hyperpolarizing current step. Current ramp stimulation protocol (top right) and voltage response of the exemplary NI neuron (bottom right) illustrating the measured properties. The AP threshold and rheobase were measured when the first AP occurred; the maximal spiking frequency was calculated using the minimal inter-spike interval. **(C, D)** Strip charts comparing membrane properties of pCCK+ (pink) and RLN3^+^ (green) NI neurons. **(E)** Pie charts illustrating the proportions of pCCK+ and RLN3+ neurons that were characterized by the occurrence and lack of sag. **(F)** Violin charts illustrating ANOVA comparison of active and passive neural properties of type I and type II NI neurons under control and MS conditions. **(G)** Current step stimulation protocol (top) and number of APs evoked in response to the stimulation (bottom). The slope of the regression was considered as the gain. (**H)** Violin charts illustrating ANOVA comparison of active neural properties of type I and type II NI neurons in control and MS conditions. **(I)** Pie charts illustrating the proportions of neurons that were characterized by the occurrence and lack of sag in control and MS rats. **(J)** Exemplary waveform of AP with comparison between type I and type II NI neurons, illustrating the measured properties. **(K)** Strip charts representing the comparison between membrane properties of pCCK+ (pink) and RLN3+ (green) NI neurons. **(L)** Violin charts representing ANOVA comparisons between type I and type II NI neurons under control and MS conditions. The following symbols were used for statistical significance determined using two-way ANOVA with a post hoc Tukey test: * MS effect, + neuronal type effect, # interaction; or ^t-tests. The number of each symbol indicates the level of statistical significance: * (p<0.05), ** (p<0.01), *** (p<0.001), **** (p < 0.0001). Please see **Supplementary Table 3, 5** and **6** for details.

Since this was the first study to characterize the electrophysiological properties of NI pCCK neurons in-depth, possible differences in membrane properties of pCCK and RLN3 NI neurons within the control group were examined. Comparison of the responses of examined neurons to incremental current steps and fast current ramp stimulation (Figure 2B) did not reveal significant differences between the active or passive membrane properties of pCCK+ and RLN3+ cells (Figure 2C-E, Supplementary Table 3), suggesting homogeneity of the electrophysiological properties of type I NI neurons, as pCCK+ and RLN3+ neurons are both type I (15). Analysis of AP properties also revealed similarities between pCCK+ and RLN3+ neurons in all but one tested feature, AHP amplitude, which was higher in RLN3 cells (Figure 2K, Supplementary Table 3). Moreover, pCCK+ and RLN3+ NI neurons did not differ in the properties of their synaptic inputs (Figure 3G, H, Supplementary Table 4).

**Figure 3.**
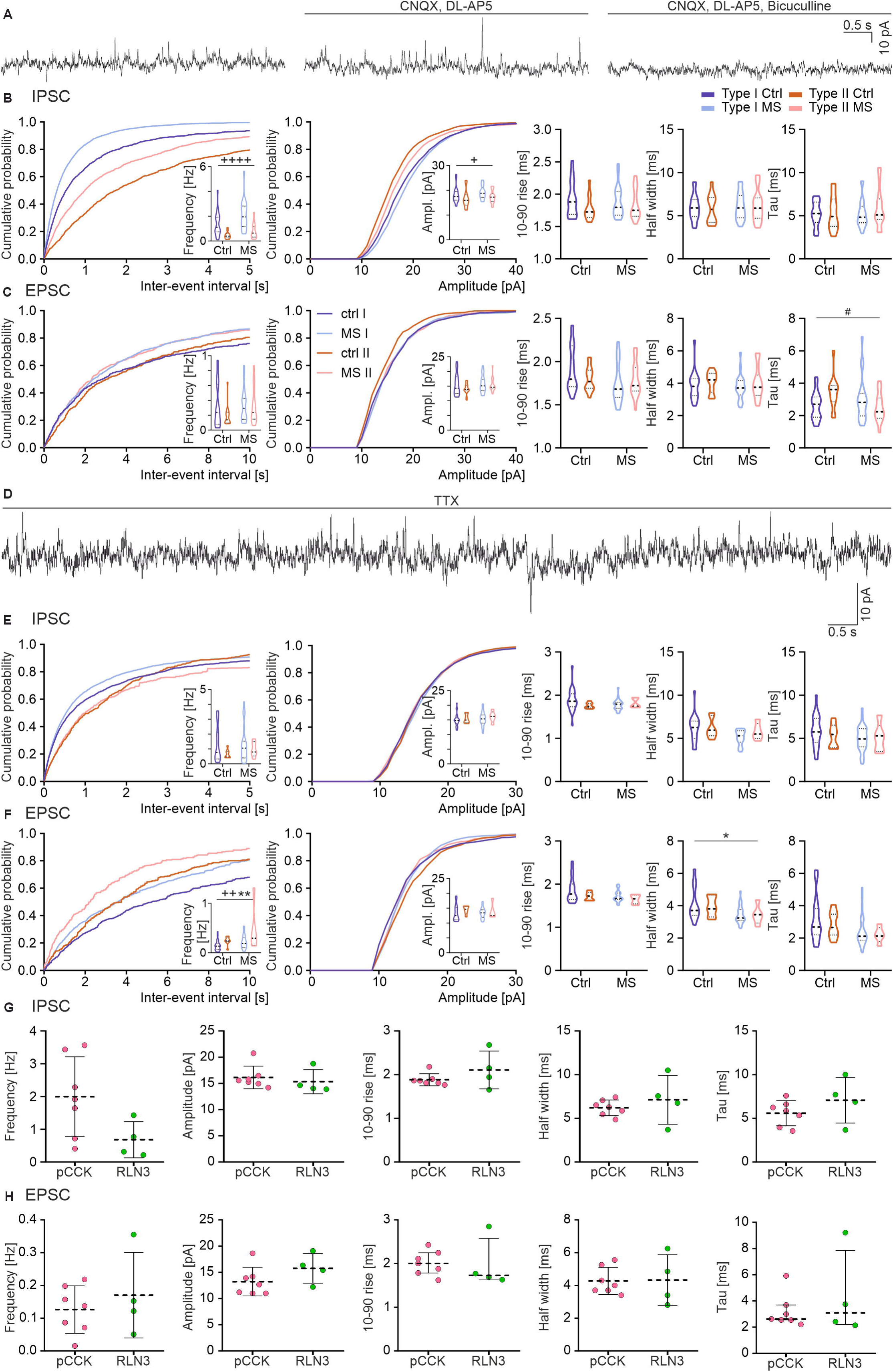
Spontaneous and miniature synaptic inputs to NI neurons. **(A)** Exemplary trace of NI spontaneous synaptic events recorded in voltage clamp at –50 mV holding potential (left), along with traces illustrating the abolition of inward currents with glutamatergic receptors antagonists (middle) and abolition of both inward and outward PSC with glutamatergic and GABAergic receptors antagonists (right). **(B)** Cumulative distribution histograms of the sIPSC (bin: 10 ms), strip chart illustrating the comparison between type I and II NI neurons in the control group and after MS (right-hand side), and strip charts representing comparison between type I and type II NI neurons in control and MS groups. **(C)** Cumulative distribution histograms of the sEPSC (bin: 10 ms), strip chart illustrating the comparison between type I and II NI neurons in the control group and after MS (right-hand side) and strip charts representing a comparison between type I and type II NI neurons in control and MS groups. **(D)** Exemplary trace of NI miniature synaptic events recorded in voltage clamp at –50 mV holding potential, with TTX added to the ACSF. **(E)** Cumulative distribution histograms of the mIPSC (bin: 10 ms), strip chart illustrating the comparison between type I and II NI neurons in the control group and after MS (right-hand side) and strip charts representing comparison between type I and type II NI neurons in the control and MS groups. **(F)** Cumulative distribution histograms of the mEPSC (bin: 10 ms), strip chart illustrating the comparison between type I and II NI neurons in the control group and after MS (right-hand side) and strip charts representing a comparison between type I and type II NI neurons in control and MS groups. **(G)** Strip charts representing a comparison between mIPSC properties of the pCCK+ (pink) and RLN3+ (green) NI neurons. **(H)** Strip charts representing comparison between mEPSC properties of the pCCK+ (pink) and RLN3+ (green) NI neurons. The following symbols were used for statistical significance determined using two-way ANOVA with a post hoc Tukey test: * MS effect, + neuronal type effect, # interaction. The number of each symbol indicates the level of statistical significance: * (p<0.05), ** (p<0.01), **** (p<0.0001). Please see **Supplementary Table 4** and **8** for details.

The possible influence of MS on the electrophysiological properties of NI neurons was examined. ANOVA and further analysis of the results revealed that in both control and MS groups, there were significant differences between type I and type II NI neurons in the majority of tested membrane properties (i.e., resistance, capacitance, action potential (AP) threshold, rheobase and neuronal gain). Importantly, MS significantly decreased the membrane capacitance in type I neurons and caused an increase in sag amplitude in type II NI neurons (Figure 2, Supplementary Figure 2, Supplementary Table 5 and 6). Additional analysis was conducted to verify whether MS affected NI neurons differently depending on their electrophysiological type (I/II), localization (NIc/NId) and neurochemical nature (i.e., presence or absence of RLN3 or pCCK). The results revealed a significant effect of MS on the sag amplitude, AP maximum frequency and neuronal gain of NIc RLN3 neurons, and on the capacitance of pCCK neurons (Supplementary Figure 2). A significant difference in sag amplitude was also observed in NIc type II neurons, that do not express either of the neuropeptides (Supplementary Figure 2). MS also influenced the time constant (tau) of type II neurons in NId and the membrane capacitance of type I neurons in NIc (Supplementary Figure 2C and D, respectively). Details of the statistical analysis are reported in Supplementary Table 7.

#### MS altered the action potential properties of NI neurons

In studies to assess the possible influence of MS on the action potential (AP) properties of NI neurons, the membrane potential of recorded cells was held at –75 mV, and a single depolarizing current pulse exceeding the excitability threshold was applied (0.75 nA, 2 ms). For the ANOVA of AHP-related properties, only the slow AHP (sAHP, occurring 25–572 ms after AP peak), present in both type I and type II NI neurons, was considered. Fast AHP (fAHP, occurring 1–9 ms after AP peak), detected only in a fraction of type II neurons, was not included in this analysis.

ANOVA revealed that type I and II NI neurons differ in terms of AP half-width, amplitude, and time to the AHP, regardless of the procedure (Ctrl/MS) to which the tested rats were subjected (Figure 2L, Supplementary Table 5). Notably, MS decreased the time to AHP in an electrophysiological type-dependent manner, particularly impacting type II neurons (Figure 2L). In-depth analysis designed to indicate the possible influence of MS on AP properties of NI neurons differing in terms of electrophysiological type, neuron localization and presence/absence of the different neuropeptides revealed specific shortening of AP (shorter AP half-width) in NIc type II neurons (Supplementary Figure 2f). Within this analysis, the fast AHP and slow AHP of type II NI neurons were considered, and notably MS caused fast AHP to occur more rapidly in type II NI neurons localized in NIc but did not affect the slow AHP (Supplementary Figure 2g). Details of the statistical analysis are reported in Supplementary Table 7.

#### MS modified synaptic activity within the NI

Synaptic activity within the NI was assessed by analyzing the 200 s long recordings in the voltage clamp configuration (holding potential –50 mV). Under these conditions, both outward and inward postsynaptic currents (PSCs) were observed, and their dependence on GABAergic and glutamatergic signaling was confirmed using respective receptor antagonists (Figure 3A), indicating that these are inhibitory PSC (IPSCs) and excitatory PSCs (EPSCs), respectively. The spontaneous postsynaptic currents (sPSCs) were recorded in standard ACSF, and frequency, amplitude, 10–90 rise, half-width and tau were evaluated for recorded inward and outward PSCs. ANOVA revealed that type I NI neurons were characterized by higher frequency, as well as higher amplitude sIPSCs than type II neurons in both examined groups (Figure 3B, Supplementary Table 8). Notably, MS impacted NI neurons in a type-dependent manner, as it produced an increase in the tau value in type I and decreased this value in type II NI neuron EPSCs (Figure 3C, Supplementary Table 8). Further analysis revealed that in the case of type II neurons, MS caused a decrease in the EPSC tau value only in NIc (Supplementary Figure 3d, Supplementary Table 9).

Miniature PSCs (mPSCs) were recorded from NI neurons in the presence of TTX in ACSF (Figure 3D). ANOVA revealed a significantly higher frequency of mEPSC in type II than in type I NI neurons in both Ctrl and MS groups (Figure 3F, Supplementary Table 8). Importantly, MS caused an increase in the frequency and a decrease in the half-width of the mEPSCs in both types of studied neurons (Figure 3F, Supplementary Table 8). Additional analysis revealed that MS increased the frequency of mEPSCs of neurons that were immunonegative for the tested neuropeptides. In positively-stained cells, MS affected synaptic activity mainly in pCCK+ neurons, as it led to a decrease in the rise time and half-width of mIPSCs, as well as a decrease in the rise time, half-width, and tau of the mEPSCs recorded from NIc pCCK+ neurons (Supplementary Fig 3g, h). MS also produced an increase in the frequency of mIPSCs in NIc RLN3+ neurons, indicating that MS influences both inhibitory and excitatory synaptic transmission in pCCK+ and only inhibitory synaptic transmission in RLN3+ neurons (Supplementary Figure 3e). Details of the statistical analysis are provided in Supplementary Table 9.

### MS produced type-dependent dendritic shrinkage of NI neurons

Following the electrophysiological experiments, dendritic tracing of biocytin-filled type I RLN3+, type I pCCK+ and type II NI neurons was performed to investigate the possible influence of MS on the morphology of these cells. On the basis of the resultant data, Sholl and additional and additional analyses of dendritic tree topological parameters were performed.

Two-way ANOVA and post-hoc tests revealed that in control rats type I pCCK+ neurons had the most complex dendritic tree among tested groups, with more intersections in its proximal part (Figure 4A, B, Supplementary Figure 4, Supplementary Table 10, 11). Interestingly, type I pCCK+ neurons were the most sensitive to MS, and MS caused significant shrinkage of their dendritic tree in its proximal and middle parts (Figure 4A, B, Supplementary Table 11). There was also a global effect, as MS exclusively lowered the maximal number of intersections and their sum in these neurons (Supplementary Figure 4, Supplementary Table 10). Concurrently, MS had no marked effect on type I RLN3 and type II cells (Figure 4A, B, Supplementary Figure 4, Supplementary Table 10, 11).

**Figure 4.**
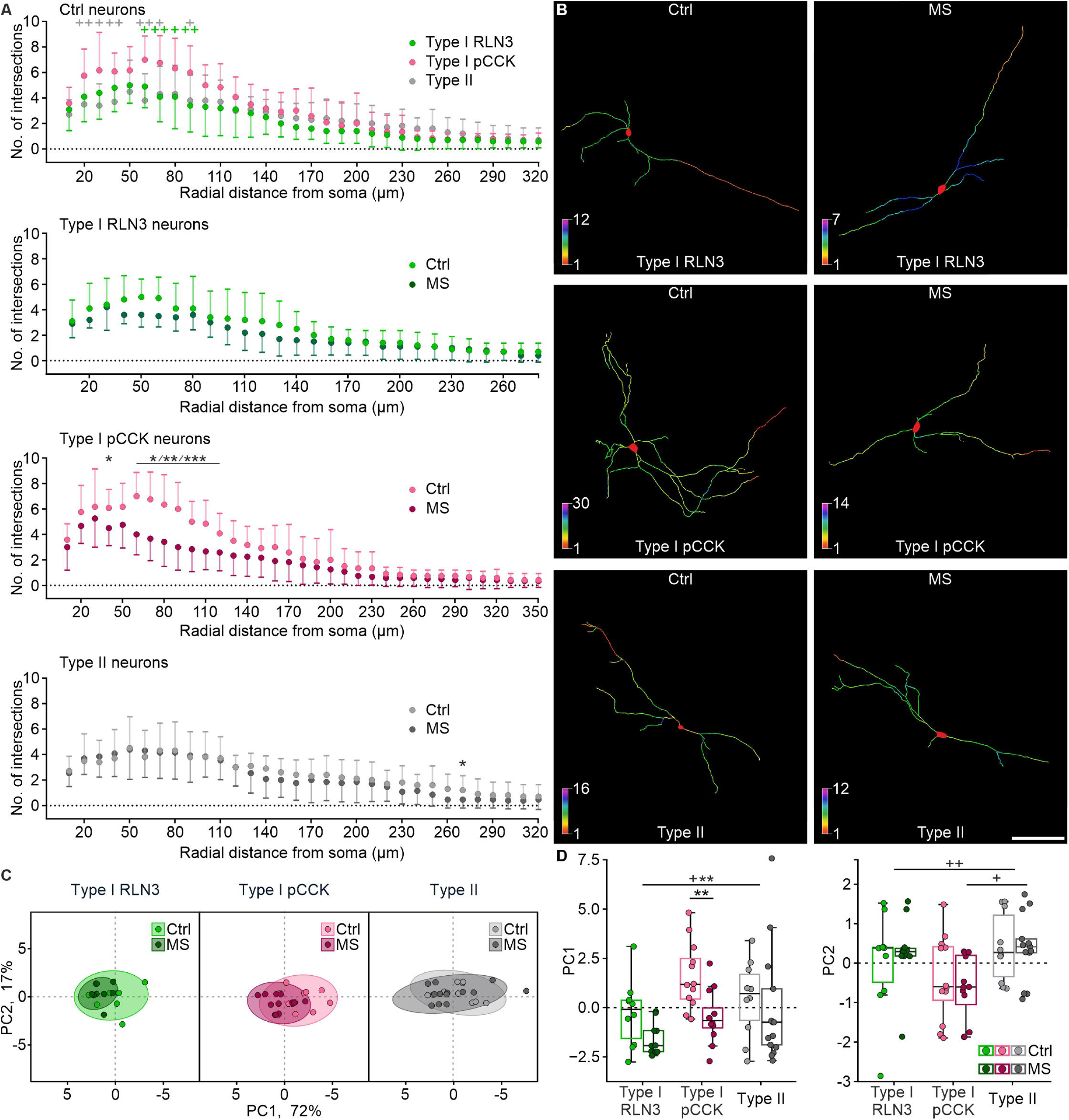
Maternal separation causes a selective dendritic shrinkage of pCCK-immunopositive NI neurons. **(A)** Sholl analysis of 3D-reconstructed control and MS NI cells, illustrating differences between the complexity of studied types of control neurons, with significantly more complex type I pCCK cells (top panel), exclusively affected by MS (lower middle panel). The x-axis range varies with the radial range of dendrites within a given cell type. **(B)** Representative skeletal 3D reconstructions of defined NI neurons from control and MS groups. Colors indicate the number of dendritic intersections with virtual Sholl spheres. Note a pronounced MS-caused dendritic atrophy of type I pCCK neurons. Scale bar = 100 μm. **(C, D)** Principal component analysis of measured topological parameters of the dendritic tree (see **Supplementary Table 12**–**14** for details). **(C)** Scatter plots of the first two principal components – a comparison of control and MS neurons of all studied types. Loadings of parameters used in the PCA are listed in **Supplementary Table 15**. **(D)** Statistical analysis of PC1 and PC2, indicating the most complex dendritic tree in the case of type I pCCK neurons and its MS-caused selective complexity decrease. Statistical significance in **(A)** and **(D)** was determined using two-way ANOVA with a post hoc Tukey test: * MS effect, + neuronal type effect. The number of each symbol indicates the level of statistical significance: * (p<0.05), ** (p<0.01), *** (p<0.001). The “+” symbol colors in **(A)** represent the difference between type I pCCK cells and type I RLN3 (green) or type II cells (gray). Please see **Supplementary Table 11** and **Supplementary Table 16** for details.

Topological parameters describing the shape of the dendritic tree, i.e., the number of (1) primary dendrites, (2) bifurcations, (3) branches and (4) dendritic tips, and (5) the total dendritic length, and (6) the maximal branch order, were analyzed using principal component analysis (PCA), in order to reduce the number of highly correlated features (Supplementary Table 12, 13). The first two principal components (PCs), explaining 89% of the observed variance (Figure 4C), were considered for further analysis. Parameters (2)–(5) were highly and positively correlated with PC1, whereas parameter (1) was negatively correlated with PC2 (Supplementary Table 14; see Supplementary Table 15 for loadings). Two-way ANOVA applied to PC1 revealed an effect of both MS and neuronal type, and a post-hoc test revealed a significant MS-induced decrease in complexity that was selective for type I pCCK neurons (Figure 4D, Supplementary Table 12, 16). In the case of PC2, a significant effect of neuronal type was observed, but post-hoc comparisons revealed significant differences only between type I pCCK and type II neuron dendritic trees in the MS group, with the former more complex than the latter (Figure 4D, Supplementary Table 12, 16).

### MS altered the expression of stress-related receptor mRNA in the NI

In order to verify the influence of MS on the expression of mRNA encoding key neuropeptides (RLN3, CCK), classical markers for inhibitory and excitatory neurons (vGAT1, vGlut2), and stress-related receptors (CRHR1, TrkA), multiplex fluorescence *in situ* hybridization was conducted using coronal sections throughout the rostrocaudal NI. Cells from two ROIs; NIc and NId, were included in the analysis (Figure 5A-C). No differences between the total number of counted control and MS neurons were observed (Supplementary Table 17).

**Figure 5.**
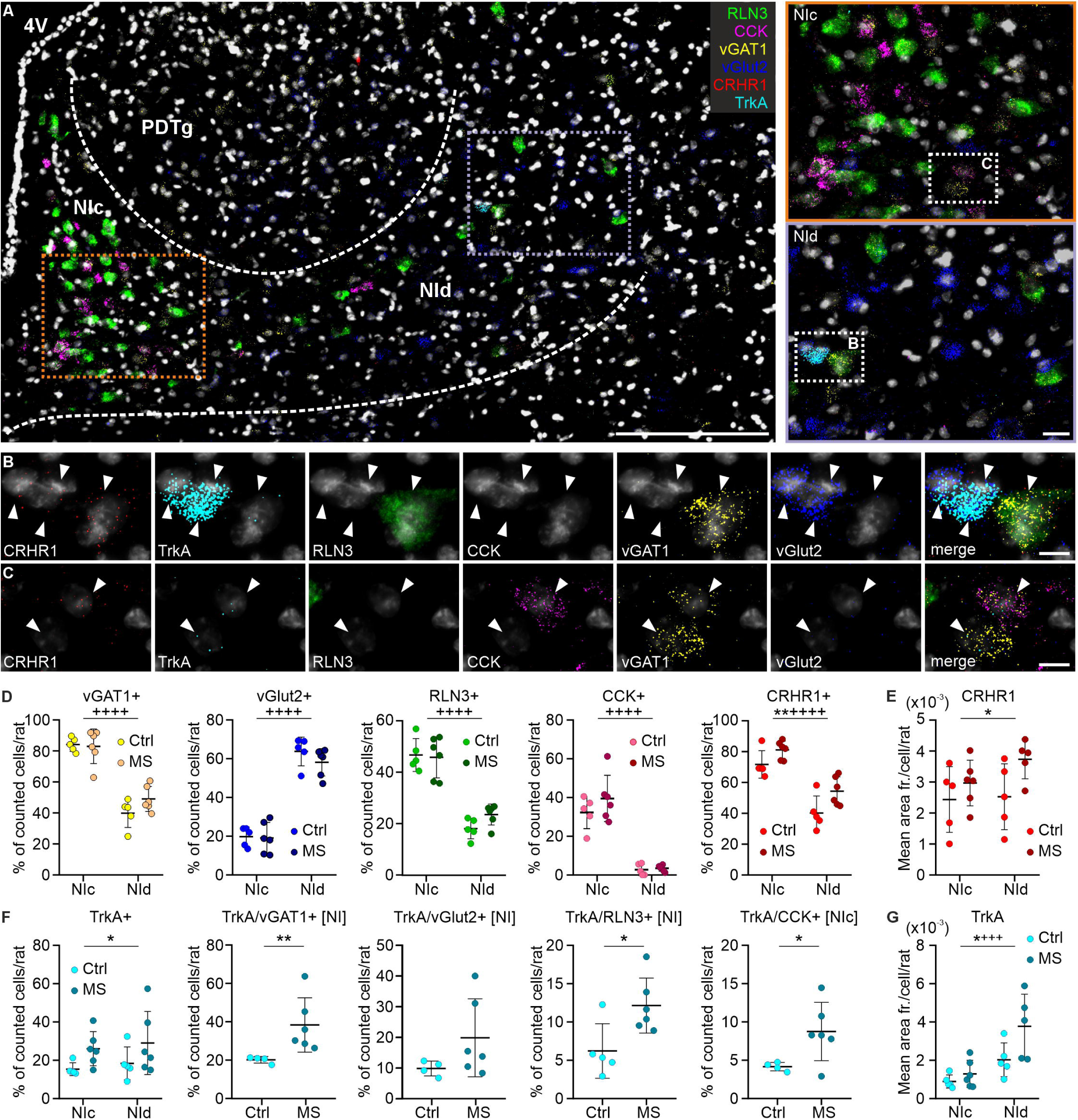
Expression of different mRNA species in control and MS rats. **(A)** Representative image of NI neurons expressing RLN3 (green), CCK (magenta), vGAT1 (yellow), vGlut2 (blue), CRHR1 (red) and TrkA (cyan) mRNAs. Scale bar: 200 μm. Enlarged sections of NIc (orange rectangle) and NId (purple rectangle) selected for cell counting are displayed on the right. Scale bar: 20 μm. **(B** and **C)** A series of images illustrating exemplary NI neurons (white arrowheads) expressing each mRNA examined with DAPI-stained nuclei (light-grey). Scale bars: 10 μm. **(D)** The percentage of counted cells expressing the examined mRNAs per rat in both groups. **(E)** The mean area fraction of CRHR1-immunofluorescent dots per cell. **(F)** The percentage of counted cells expressing TrkA mRNA in control and MS rats, and TrkA in combination with other studied mRNA species. **(G)** The mean area fraction of TrkA-immunofluorescent dots calculated per single NI cell. Statistical significance was determined by two-way ANOVA: * MS effect, + localization effect; * t test (with Welch correction if needed) or Mann-Whitney test. The number of each symbol indicates the level of statistical significance: * (p<0.05), ** (p<0.01), *** (p<0.001), **** (p<0.0001). Please see **Supplementary Table 17-22** for details. Abbreviations: 4V, 4th ventricle; NIc, nucleus incertus pars compacta; NId, nucleus incertus pars dissipata; PDTg, posterodorsal tegmental nucleus; RLN3, relaxin-3; CCK, cholecystokinin; CRHR1, corticotropin-releasing hormone receptor type 1; TrkA, tropomyosin receptor kinase A; vGAT1, vesicular γ-aminobutyric acid (GABA) transporter; vGlut2, vesicular glutamate transporter.

Counting of RLN3-mRNA-positive (Rln3+) and CCK-mRNA-positive (Cck+) cells, followed by two-way ANOVA revealed a significant NIc/NId localization effect, without an MS effect (Figure 5D, Supplementary Table 18). A similar effect was observed for NI neurons expressing vGAT1 mRNA (vGAT1+) and vGlut2 mRNA (vGlut2+) (Figure 5D, Supplementary Table 18). Importantly, in addition to the NIc/NId localization effect, a significant MS effect was observed for the NI cell population expressing CRHR1 mRNA (CRHR1+), consisting of various species of NI neurons, including vGAT1+, vGlut2+, Rln3+ and Cck+ cells (Figure 5B, C and Supplementary Table 19). The percentage of CRHR1+ cells was greater in NIc under control conditions and MS increased the percentage in both NI zones (Figure 5D, Supplementary Table 18). Moreover, in rats subjected to MS, an increase in the level of CRHR1 mRNA within individual NI neurons was observed, as the mean area fraction of CRHR1 mRNA-related immunofluorescent dots per cell was higher in MS rats (Figure 5E, Supplementary Table 20).

MS also produced an increase in the number of NI cells expressing TrkA mRNA (TrkA+, Supplementary Table 18), with vGAT1+, vGlut2+, Rln3+ and Cck+ cells observed (Figure 5B, C). The increase in the number of TrkA+/vGAT1+ and TrkA+/Rln3+ neurons was observed in the whole NI area, and the number of TrkA+/Cck+ cells increased specifically in NIc (the main area of CCK synthesis; Figure 5F, Supplementary Table 21). Notably, the mean area fraction of TrkA mRNA-related immunofluorescent dots calculated per individual NI neuron was also increased in MS rats and generally higher in NId (Figure 5G, Supplementary Table 20).

## DISCUSSION

In the current study we demonstrated that the NI, an important element of the central stress response system implicated in alcohol seeking, is highly sensitive to ELS. Specifically, we observed a decrease in the basal activity of NI RLN3 neurons in adult male rats exposed to ELS in the form of MS between PND2–14. Additionally, we observed that ELS leads to an increase in the number of CCK-positive neurons within the NI, coinciding with a change in their acute stress responsiveness in adulthood. Concurrently, we observed that ELS produced a severe impairment in CCK neuron dendritic tree structure. We also demonstrated that ELS modulated the electrophysiological membrane properties, and synaptic transmission of NI neurons, in a cell-type-specific manner. Our data also revealed that ELS increased levels of mRNA encoding CRHR1 and TrkA mRNAs in the NI of adult male rats.

### ELS and stress-responsiveness of NI neurons

NI consists mainly of GABAergic projection neurons that innervate a wide range of forebrain structures (51,52), and has been implicated in the control of contextual memory, arousal and responses to stress (22,23,27,53). Here we demonstrated that NI neurons are sensitive to ELS, and that early-life disturbances may alter the functioning of NI RLN3 and CCK neurons. Notably, we revealed that both these neuronal populations reacted to acute restraint stress with elevated activation, but ELS affected them differently. ELS disrupted the proportions of active and inactive NI CCK neurons (c-Fos protein positive and negative, respectively) and significantly increased the number of non-activated CCK neurons. These findings suggest ELS leads to dysregulation of the adult NI CCK neural network, including changes in its stress reactivity. As the current study is the first to directly link this NI cell type with neurogenic stress, further investigations are needed to elucidate the underlying mechanisms (possibly involving CRHR1 and TrkA actions) and the precise functional and behavioral implications. Unlike CCK NI neurons, RLN3 NI neurons have been proven reactive to various stressors in rats, including acute restraint, as indicated by elevated c-Fos expression (35,54). Furthermore, both c-Fos expression and electrical activity were reported to be elevated in RLN3 NI neurons after intracerebroventricular (ICV) CRH injection (27,35), and CRH excitatory action in the NI was blocked by a CRHR1 antagonist under *ex vivo* conditions (27), indicating a direct role of CRHR1 in CRH action in the NI. Our current results are consistent with these reports and confirm activation of NI neurons by acute stress. Concurrently, we demonstrated that ELS decreased the number of activated RLN3 NI neurons in adult brain, independently of the restraint stress condition, which may underpin their dampened stress-responsiveness.

### ELS-related functional and structural alternations in the NI: possible mechanisms and consequences

We demonstrated that while the total number of neurons expressing RLN3 and CCK (mRNA and peptide) was not significantly changed by ELS, the electrophysiological properties of these cells were altered by ELS experience. The RLN3 and CCK NI neurons are mostly separate populations, but are both type I NI cells (15), and our current data indicate that the passive and active membrane properties of RLN3 and CCK NI neurons are very similar. Importantly, we observed that ELS influenced the physiology of NI neurons in a specific fashion. While most of the passive and active membrane properties of NI neurons remained unchanged in adult rats exposed to ELS, the amplitude of the voltage sag, a characteristic of activating the hyperpolarization-activated cyclic nucleotide-gated (HCN) channel current (Ih), was significantly altered in an electrophysiological– and neurochemical-dependent manner. We observed that MS significantly reduced the sag amplitude in type I RLN3 NI neurons, but increased it in type II neurons. Activation of HCN channels can exert dual effects, either excitatory or inhibitory, depending on the cell type (55,56). It can lead to membrane depolarization and an increase in neuronal excitability or, by increasing membrane conductance, to attenuation of membrane potential alternations, reduction of sensitivity to incoming stimuli and a decrease in postsynaptic currents summation (57). In the current study, we observed that the sag amplitude in RLN3 NI neurons was significantly decreased by ELS experience, which was accompanied by an increase in neuronal gain in these cells. This indicates a possible ELS-induced decrease in the expression of HCN channels (or a lower level of their activation) in RLN3 neurons, with a consequent enhancement of the synaptic input summation capacity, increased excitability, and possible increased reactivity to stress-related stimuli. Considering this, the ELS-related decreased activation level might act as an adaptive mechanism to mitigate this effect. At the same time, in NIc type II neurons, ELS caused a significant increase in the sag amplitude, that was accompanied by an increase in the neuronal gain. These data, together with the ELS-induced increase in maximal firing frequency and shortened duration of the action potential, indicate a possible higher responsiveness of NIc type II neurons to incoming salient stimuli, in adult rats exposed to maltreatment during development. Notably, our previous data indicated that type I and II NI neurons belong to different neuronal circuits (15,20), and here we demonstrated the ELS sensitivity of both neuronal types, which strongly implies the potential adverse impact of maltreatment during early life stages on distinct neuronal systems, that are under NI control.

The alterations in the electrophysiological properties of NI neurons induced by ELS, were accompanied by selective and pronounced dendritic shrinkage of type I CCK NI neurons. It has been previously demonstrated that ELS can dramatically alter neuronal morphology, and rodents experiencing ELS exhibit decreased dendritic arborization in the prefrontal cortex and hippocampus (58,59), and this detrimental influence is accompanied by modification of synaptic signaling (58,60). In accordance with these data, the ELS-induced shrinkage of CCK neuron dendritic trees was accompanied by significantly altered kinetics of both excitatory and inhibitory synaptic currents, indicating a postsynaptic origin of the observed changes. These findings reflect important neuroanatomical and functional consequences of ELS on the NI, which may underlie behavioral impairments related to NI-controlled functions, including reduced spatial memory, enhanced threat learning, and the increased vulnerability to substance abuse observed in humans and animals who have experienced this kind of stress (6,61,62).

### The possible role of NI CCK neurons in stress response

Very little is known about the precise function of NI CCK neurons, which we have shown are highly sensitive to ELS. Notably, an early study of CCK neurons that were described as located in the “caudal dorsal raphe nucleus” were shown to influence behavioral components of stress-adaptation responses and to impact the HPA axis response to chronic stress (63). However, closer examination revealed that these neurons were in fact NI CCK neurons, and in our recent report we demonstrated that, together with RLN3 neurons, NI CCK cells form the NI-MS pathway (15), which implies their possible involvement in theta rhythm, spatial navigation, arousal and motivation-related behavior control (23,64–67).

Importantly, existing research has demonstrated a substantial involvement of CCK neurons in the regulation of substance abuse behavior, including alcohol abuse, through their heightened sensitivity to various stress factors, including ELS (45–47). Similarly, several studies have demonstrated that the NI and RLN3/RXFP3 system contribute to alcohol preference in rodents (27,37). Pretreatment with CRHR1 antagonists reduced alcohol seeking in mice and rats (68,69), and CRHR1 signaling within the NI was strongly implicated in this process, as intra-NI CRHR1 antagonist injections strongly attenuated stress-induced reinstatement of alcohol seeking (30).

Numerous studies also report an increase in drug abuse vulnerability in adults who experienced ELS (6). In the current study, ELS significantly increased the number of NI CRHR1 mRNA-expressing neurons and the level of mRNA expression per cell, in accordance with reports that ELS disrupts the normal development of the CRH/CRHR1 system, and leads to a net reinforcement of HPA axis activity in humans and rodents (40). Thus, current data suggest the possible involvement of ELS-induced changes in NI CCK neuron anatomy and physiology, as well as CRHR1 expression in the NI, in the increased susceptibility to substances of abuse in adult individuals who have experienced childhood neglect.

### Implications of ELS-induced TrkA mRNA dynamics in NI neurons

We also demonstrated that ELS increased the number of NI TrkA mRNA-expressing cells, and the level of TrkA mRNA per neuron in the NI. Although a previous study reported that a promoter region for TrkA was selectively co-expressed with RLN3 in NI neurons (50), other gene promoters were not tested, and here we demonstrated that TrkA mRNA is expressed by GABA/RLN3 and GABA/CCK neurons, as well as glutamate NI neurons. Notably, we observed that ELS-induced upregulation of TrkA mRNA occurred only in GABA, and not glutamate NI neurons, indicating a possible functional distinction between these neuronal subpopulations. Indeed, a recent study reported that in pontine central gray (PCG), of which NI is an integral part, GABA and glutamate neurons encode positive and negative valence, respectively, and that activation of these two populations resulted in preference (GABA) and avoidance behavior (glutamate) (70). Our findings indicate that NI GABAergic and glutamatergic neuronal populations exhibit differential susceptibility to ELS, which may contribute to the atypical processing of valence-related information, a phenomenon documented in subjects with a history of ELS (71).

Several studies have shown that alcohol dependence is associated with alterations in plasma levels of NGF (72) and after alcohol withdrawal a rapid increase in the serum NGF concentration was observed (73–75). It was also shown that ELS causes an increase in the level of NGF in the hippocampus, cerebral cortex and hypothalamus (76,77), leading to impaired synaptic plasticity and behavioral deficits (78). These data suggest that anxiety resulting from both ELS and alcohol use and withdrawal triggers NGF release. Notably, NGF increases TrkA expression (79), and our current data indicate that TrkA mRNA in the NI is increased by ELS, which together with an increase in CRHR1 mRNA expression, may translate into an altered sensitivity of the NI to stress in adulthood. Moreover, the increased basal expression of TrkA resulting from ELS may contribute to the augmentation of alcohol withdrawal-induced anxiety and promote stress-induced relapse of its consumption.

Additionally, ELS-induced upregulation of TrkA mRNA expression may be associated with enhanced protective mechanisms within the stress-sensitive NI neurons, as an established function of NGF is to facilitate neuronal survival (80). However, the maladaptive or recurrent activation of NGF synthesis during the early postnatal period may impact stress sensitivity in adulthood and heighten susceptibility to stress-related psychopathology (43).

### Concluding remarks

Collectively, current findings advance our understanding of the neural mechanisms underlying ELS-related neuronal impairments, and together with previous data indicating the NI as a structure critically involved in contextual memory control (22) and stress-induced reinstatement of alcohol seeking (30,31,81), position the NI as a possible node linking ELS with compromised stress responses and greater vulnerability to substance abuse.

## ACKNOWLEDGEMENTS AND DISCLOSURES

This work was supported by the National Science Centre, Poland – UMO-2017/27/N/NZ4/01545 (AG), UMO-2023/49/B/NZ4/01885 (AB), and UMO-2021/41/N/NZ4/04499 (SD) and by the European Union’s Horizon Europe research and innovation program under Marie Sklodowska-Curie grant agreement No 101086247 – PsyCoMed Project (AB and ALG).

We thank Patrycjusz Nowik for his excellent animal husbandry and Aleksandra Celary for her help with the maternal separation procedure.

A.G. and A.B. designed the experiments. A.G. and P.S. performed the maternal separation procedure. A.G., P.S. and A.T. conducted the c-Fos immunohistochemistry experiments. P.S. conducted the electrophysiological experiments and analyzed the resultant data. A.G. and A.T. conducted the in situ hybridization experiments. A.G., A.T., S.D. and A.N. analyzed the resultant microscopy data. Z.S. performed the statistical analysis. A.B., A.G., P.S. and A.L.G. drafted and edited the article, and all authors provided comments and corrections.

The authors report no conflicts of interest.

