## Supplementary Materials and Methods for "Early-life adversity alters adult nucleus incertus neurons: implications for neuronal mechanisms of increased stress and compulsive behavior vulnerability"

#### Animals and treatment

Male Sprague-Dawley rats (control, Ctrl, n = 33, from 8 litters; and maternally separated, MS, n = 39, from 9 litters) from the Institute of Zoology and Biomedical Research animal facility of the Jagiellonian University in Krakow, Poland, were used in the described experiments. The rats were housed with food and water available *ad libitum*, on a standard light/dark cycle (lights on 08.00–20.00), temperature 22-23°C. The experiments were approved by the 2nd Local Institutional Animal Care and Use Committee (Krakow, Poland) and conducted in accordance with the EU Directive 2010/63/EU on the protection of animals used for scientific purposes. All efforts were made to minimize the number of animals used and their suffering.

*Maternal separation procedure*

Pregnant dams, housed as described, were selected and isolated from males to avoid reimpregnation and bearing a second litter during the maternal separation (MS) procedure. The litters were standardized to a fixed number of 5–9 male pups. The day of birth was defined as postnatal day 0 (PND 0), and the MS procedure was conducted during PND 2–14, as described (1). Briefly, the dams were moved to separate holding cages for 3 h daily (Zeitgeber time (ZT) 3–6), leaving the litters alone as a group in their home cage. On PND 28 the rats were weaned, transferred to separate cages, and housed as family groups until further experimental procedures.

#### Acute stress and c-Fos expression in neurochemically-defined nucleus incertus neurons

*Restraint stress procedure and brain fixation*

On PND 55–56, control (Ctrl) and maternally separated (MS) rats were subjected to 30 min restraint stress in the experimental room (control stressed, Ctrl S, n = 5; MS stressed, MS S, n = 6), to determine the influence of MS on stress-induced c-Fos protein expression in the NI. Briefly, the rats were restrained for 30 min in custom-made disposable transparent plastic bag restrainers with a breathing hole at the tip, during ZT 1–5. As a control condition, rats from Ctrl (n = 6, Ctrl NS) and MS (n = 5, MS NS) groups were kept in transport cages in the experimental room for 30 min without restraining (rats from the other groups were restrained on a different day). At 1 h (sufficient time for c-Fos protein expression; (2)) after the end of the restraint procedure (Ctrl S and MS S groups) or after the end of the exposure to the experimental room (Ctrl NS and MS NS groups), rats from all groups were pre-anesthetized with isoflurane (Aerrane, Baxter, Poland) and deeply anesthetized with pentobarbital (Morbital, Biowet, Poland; intraperitoneally, dose: 2 ml/kg). Subsequently, rats were transcardially perfused with phosphate-buffered saline (PBS, 4°C, pH 7.4, ~250 ml) followed by 4% phosphate-buffered formaldehyde solution (4°C, pH 7.4, ~250 ml). At the end of the procedure, brains were collected and postfixed overnight (4% formaldehyde, 4°C).

*Immunohistochemical staining, imaging and cell counting*

After post-fixation, brains were washed in PBS and transferred to 30% sucrose solution in PBS (4°C, ~24 h). Next, the brainstem area containing the NI was cut into 45 µm coronal sections on a freezing microtome. Every fourth section was incubated in 10% normal donkey serum (NDS, Jackson ImmunoResearch, West Grove, PA, USA) and 0.3% Triton X-100 (Sigma-Aldrich, cat. no. X100) PBS solution (1 h, RT). Sections were then transferred into a solution containing primary antibodies against relaxin-3 (RLN3, mouse monoclonal, 1:10, provided by The Florey Institute of Neuroscience and Mental Health, Parkville, Australia) and c-Fos (rabbit polyclonal, 1:10000, Abcam, Cambridge, UK, cat. no. ab190289), 2% NDS and 0.3% Triton X-100 in PBS, and incubated for ~72 h at 4°C. After washing in PBS, sections were placed in a secondary antibody solution, containing Alexa Fluor 488 anti-mouse (donkey, 1:400, Jackson ImmunoResearch, cat. no. 715-546-150), Cy3 anti-rabbit (donkey, 1:400, Jackson ImmunoResearch, cat. no. 711-165-152) and 2% NSD in PBS at 4°C, overnight. Sections were then washed and all described steps were repeated, with a primary antibody against pro-cholecystokinin (pCCK, rabbit polyclonal, 1:200, Frontier Institute Co., Ltd, Hokkaido, Japan, cat. no. CCK-pro-RB-Af350). After washing in PBS, sections were incubated with Alexa Fluor 647 anti-rabbit secondary antibody (donkey, 1:400, Jackson ImmunoResearch, cat. no. 711-606-152; overnight, 4°C) and mounted on glass slides using Fluoroshield with DAPI (Sigma Aldrich, cat# F6057).

Z-stack panoramic images of matching brainstem sections containing NI (three per brain) were collected using Axio Imager M2 fluorescent microscope (Zeiss, Gottingen, Germany), equipped with an automatic z-stage, Axiocam 503 mono camera (Zeiss) EC Plan-Neofluar 20x/0.50 M27 objective (scaling: 0.227 × 0.227 × 1.380 µm/pixel, respectively in the x, y and z axis). To improve the signal-to-noise ratio, the images were subsequently postprocessed with Zen 3.1 blue edition (Zeiss) and ImageJ software (3).

Neurons immunopositive for RLN3 (RLN3+) and pro-CCK (pCCK+), co-expressing c-Fos protein (c-Fos+) or without such co-expression (c-Fos–), were counted using an ImageJ Cell Counter plugin. Since the rat NI has been anatomically divided into the pars compacta (NIc) and pars dissipata (NId) (4), cells in the NIc and NId were counted separately. Neurons immunopositive for both RLN3 and pCCK were excluded from the analysis due to the rarity of their occurrence.

**Electrophysiological patch clamp recordings**

A total of 38 male rats (5–8 weeks old, Ctrl group; n = 17, MS group; n = 21) were used in electrophysiological experiments. Rats were deeply anesthetized with isoflurane (Aerrane, Baxter, Poland) and decapitated (ZT 1–2). Brains were dissected from the skull in carbogenated, ice-cold artificial cerebrospinal fluid (ACSF), containing (in mM): 185 sucrose, 25 $NaHCO_3$, 3 KCl, 1.2 $NaH_2PO_4$, 2 $CaCl_2$, 10 $MgSO_4$ and 10 glucose, pH 7.4; osmolality 290–300 mOsmol/kg) and cut into 250 µm coronal slices on a Leica VT 1000S vibrating microtome (Leica Instruments, Heidelberg, Germany). NI containing slices were subsequently transferred to an incubation chamber containing warm (32°C, heating was turned off after transfer of the first slice), carbogenated standard ACSF (sACSF) containing (in mM): 118 NaCl, 25 $NaHCO_3$, 3 KCl, 1.2 $NaH_2PO_4$, 2 $CaCl_2$, 2 $MgSO_4$, and 10 glucose (pH 7.4; osmolality 290–300 mOsmol/kg).

After at least 90 min of recovery, slices were transferred to the recording chamber, with constant perfusion (2 ml/min) of fresh, warm (32°C), carbogenated sACSF. The recording chamber was mounted on the fixed stage of an Axioskope 2 microscope (Zeiss). Whole-cell configuration was obtained under visual control with negative pressure obtained by pressure controller (ez-gSEAL 100B, NeoBiosytems, San José, CA, USA), delivered through recordings micropipettes of resistance between 5 and 8 MΩ. Recording micropipettes were pulled on a horizontal puller (P-1000, Sutter Instruments, Novato, CA, USA) using borosilicate glass capillaries (Sutter Instruments). The pipette solution contained (in mM): 125 potassium gluconate, 2 $MgCl_2$, 4 $Na_2ATP$, 0.4 $Na_3GTP$, 5 EGTA, 10 HEPES (pH 7.3; osmolality 290–300 mOsmol/kg) and biocytin (0.05%) which allowed for subsequent immunofluorescent identification of recorded neurons. The calculated liquid junction potential for sACSF and the intrapipette solution was +15 mV, and this value was subtracted from the data. All reagents used for the ACSF and intrapipette solution were purchased from Sigma-Aldrich (Darmstadt, Germany). An SEC 05LX amplifier (NPI, Tamm, Germany) and Signal and Spike2 (Cambridge Electronic Design, Cambridge, UK) software were used for data acquisition and further analysis (low-pass filtered at 3 kHz and digitized at 20 kHz), as well as a custom-made script in MATLAB (MathWorks Inc., Natick, MA, USA). All drugs were delivered via a bath perfusion system.

The experimental protocols used in voltage-clamp conditions (holding voltage –75 mV) were as follows; (a) voltage ramp (1.3 s, –120 mV to +20 mV), (b) voltage steps (0.5 s, –120 mV to 0 mV, 10 mV increments), (c) test for potassium A current (hyperpolarization to –100 mV for 0.5 s and subsequent 0.5 s voltage steps from –75 mV to 25 mV, 10 mV increments); and in current-clamp conditions (membrane potential set at –75 mV); current ramp (1 s, 0 nA to 1 nA), current steps (0.5 s, –150 pA to +150 pA, 10 pA increments), single pulse to evoke action potential (2 ms, 0.75 nA). Next, the spontaneous activity of the recorded neurons was examined under current-clamp conditions (0 pA holding current) for at least 300 s, and subsequently spontaneous postsynaptic currents (sPSCs) were recorded under voltage-clamp conditions (holding potential –50 mV), for at least 300 s. Calculated reversal potential for Cl, Na and K currents in our patch-clamp recordings equaled –90.46, +72.33 and –98.02 mV respectively; therefore, at the –50mV holding potential, outward events represented IPSCs, whereas inward events represented EPSCs.

For examination of the miniature postsynaptic currents (mPSCs), a separate group of neurons was recorded in the presence of tetrodotoxin (TTX, 0.5 μM, Tocris Bioscience, Cat No. 1069, Bristol, UK) in ACSF. This group underwent the same tests in voltage- and current-clamp protocols. Both sPSCs and mPSCs were detected offline and analyzed using Mini Analysis software (Synaptosoft Inc., Fort Lee, NJ, USA) and were performed by experimenters blinded to the treatment conditions.

*Post-recording immunostaining*

After recording, brain slices were transferred to 4% formaldehyde in PBS (24 h, 4°C). Slices were washed in PBS and incubated in PBS solution containing 10% NDS, 0.6% Triton X-100 (overnight, 4°C). Next, after another washing in PBS, the slices were incubated in PBS solution containing primary antibodies against RLN3 (1:25) and pCCK (1:200), ExtrAvidin-Cy3 (biocytin binding protein, 1:200, Sigma-Aldrich, cat. no. E4142), 2% NDS and 0.3% Triton X-100 for 48–72 h at 4°C. Subsequently, slices were washed in PBS and incubated in PBS solution containing 2% NDS, Alexa Fluor 647-conjugated donkey anti-rabbit (1:400, Jackson ImmunoResearch, cat. no. 711-605-152, West Grove, PA, USA) and Alexa Fluor 488-conjugated donkey anti-mouse secondary antibodies (1:400, 24 h, 4°C). After a final washing, the slices were mounted on glass slides and coverslipped with Fluoroshield with DAPI.

### Dendritic tracing and morphological analysis

After patch-clamp recordings, the influence of MS on the dendritic tree morphology of different types of NI neurons was examined in electrophysiological type I RLN3-immunopositive (control, n = 10, from 5 rats, and MS, n = 10, from 9 rats) and pro-CCK-immunopositive (pCCK, control, n = 12, from 8 rats, and MS, n = 12, from 7 rats) neurons, and in electrophysiological type II cells (control, n = 10, from 6 rats, and MS, n = 13, from 6 rats) that were well-stained against biocytin, with clearly visible dendritic trees and with no evident truncations. Selected NI cells were imaged using a confocal laser scanning microscope (LSM 780 on Axio Observer Z1, Zeiss) with a Plan-Apochromat 20×/0.8 M27 objective (scaling: 0.17 × 0.17 × 1.80 µm/pixel, respectively in the x, y and z axis) and subjected to semi-automatic dendritic tracing using Simple Neurite Tracer plugin in ImageJ and saved as *.swc files for further analysis (5). Simultaneously a built-in function was used to conduct 3D Sholl analysis (6), with a 10 µm step size between virtual spheres intersecting with dendrites. The number of intersections per single radius, the sum of intersections and the maximum number of intersections per radius in each studied group of cells were analyzed. Finally, L-Measure software (7) was used to obtain topological parameters of the dendritic tree.

### Multiplex fluorescent *in situ* hybridization

Multiplex fluorescent *in situ* hybridization (RNAscope™ HiPlex Assay for AF488, Atto550 and Atto647 detection, Advanced Cell Diagnostics (ACD), Hayward, CA, USA) was performed using 16 µm fresh frozen brain sections from 5 control and 7 MS rats (PND 51–57; 3 matched NI containing slices/brain; brains collected during ZT 4–7). All procedures were performed following the user manual provided by the manufacturer and as described (8). The following probes were applied for round 1 (R1): RLN3 (Rn-Rln3-T1, cat. no. 1037211-T1, ACD), CCK (Rn-Cck-T2, cat. no. 532851-T2, ACD), vGAT1 (Rn-Slc32a1-T3, cat. no. 424541-T3, ACD), and round 2 (R2): CRHR1 (Rn-Crhr1-T4, cat. no. 318911-T4, ACD), vGlut2 (Rn-Slc17a6-T5, cat. no. 31701-T5, ACD) and TrkA (Rn-Ntrk1-T6, cat. no. 402611-T6, ACD).

Images of an ROI (1936 × 1460 pixels/220 × 166 µm, the same area for R1 and R2) located in NIc and NId were acquired for each slice, using an Axio Imager M2 fluorescent microscope (Zeiss) with an automatic z-stage, Axiocam 503 mono camera (Zeiss) and an EC Plan-Neofluar 40×/1.3 Oil M27 objective (scaling: 0.114 × 0.114 × 0.280 µm/pixel, respectively in the x, y and z axis). The images were postprocessed using Zen (3.3 blue edition and 2.3 SP1 black edition, Zeiss), ImageJ and HiPlex Image Registration Software v1.0 (ACD), and cells were counted with an ImageJ Cell Counter plugin. If at least two unambiguous dots of specific fluorescence were detected within a cell boundary (identified by the presence of the DAPI-stained nucleus and/or a cell-shaped distribution of fluorescent mRNA signal), it was classified as expressing a specific mRNA molecule. A custom-made ImageJ macro was applied to estimate the area fraction of the ROI occupied by fluorescent dots representing CRHR1 and TrkA mRNA.

### Statistical analysis

Cell counting, morphological and postsynaptic currents analyses were performed by experimenters blinded to the treatment conditions. The data were analyzed using Prism 8

(GraphPad Software, USA) and R (R Foundation for Statistical Computing, Vienna, Austria) software. The normality of the data distribution and the homogeneity of variance were verified with the Shapiro–Wilk test and the Levene test. Data points identified as outliers using a ROUT test (Q=1%) were excluded from the analysis. Data meeting the criteria for the use of parametric tests were analyzed using unpaired t test (with Welch's correction, if the incorporated F test p value <0.05) or type III two-way ANOVA. Remaining data were analyzed using a Mann-Whitney test or type III two-way ANOVA with White's correction. After two-way ANOVA Tukey's test (Prism) or emmeans function with Tukey adjustment method was used for post-hoc comparisons (R emmeans package), whenever necessary. Differences in the sag frequencies and proportions of c-Fos+ and c-Fos– neurons between groups were analyzed using Fisher's exact test. All data in graphs and tables are presented as mean ± SD or median ± IQR, depending on the tests used.
