## Supplementary Figure Legends for "Early-life adversity alters adult nucleus incertus neurons: implications for neuronal mechanisms of increased stress and compulsive behavior vulnerability"

**Supplementary Figure 1. Influence of restraint stress and MS on c-Fos expression in different NI areas**. **(a)** The number of all NI and NId c-Fos+ cells was increased by restraint stress. **(b)** The number of RLN3+ NIc but not NId c-Fos+ cells was increased by restraint stress. (**c)** The number of RLN3+ NIc but not NId c-Fos- cells was decreased by restraint stress. **(d)** Restraint stress and MS did not affect the number of all counted RLN3+ NI and NId neurons. **(e)** Restraint stress and MS differentially impacted the number of all counted pCCK+ NIc neurons. Statistical significance was determined using two-way ANOVA: $ restraint stress effect, # interaction of MS and restraint stress. The number of each symbol indicates the level of statistical significance: * ($p<0.05$), ** ($p<0.01$). Please see **Supplementary Table 1** for details. Abbreviations: NI, nucleus incertus; NIc, nucleus incertus pars compacta; NId, nucleus incertus pars dissipata; NS, not subjected to restraint stress; MS, maternal separation; pCCK, pro-cholecystokinin; RLN3, relaxin-3; S, subjected to restraint stress.

**Supplementary Figure 2. Impact of MS on the properties of NI neurons**. **(a)** The neuronal types compared. (**b)** Voltage step stimulation protocol (left) and current responses of exemplary NI neurons, illustrating the occurrence of type A potassium current in type I NI neuron (middle), and lack of type A potassium current in type II NI neuron (right). **(c-g)** Violin graphs comparing the membrane and spike properties of control and MS groups. Asterisks represent statistical significance: * ($p<0.05$), ** ($p<0.01$), *** ($p<0.001$), **** ($p<0.0001$). Please see **Supplementary Table 7** for details.

**Supplementary Figure 3. Impact of MS on the properties of the neural input to NI neurons**. **(a, b)** Violin graphs comparing properties of outward spontaneous postsynaptic currents (sIPSCs) in control and MS groups. **(c, d)** Violin graphs comparing properties of inward spontaneous postsynaptic currents (sEPSCs) in control and MS groups. **(e, f)** Violin graphs comparing the properties of outward miniature postsynaptic currents (mIPSC) in control and MS groups. **(g, h)** Violin graphs comparing the properties of inward miniature postsynaptic currents (mEPSCs) in control and MS groups. Asterisks represents statistical significance: * ($p<0.05$), ** ($p<0.01$), *** ($p<0.001$), **** ($p<0.0001$). Please see **Supplementary Table 9** for details.

**Supplementary Figure 4. Maternal separation selectively decreased the complexity of pCCK-positive NI neurons**. **(a**, **b)** Additional parameters of the Sholl analysis of defined control and MS NI neurons: **(a)** sum of dendritic intersections with Sholl spheres; **(b)** maximal number of intersections per single sphere radius. Note the significantly higher complexity of type I pCCK-positive neurons, compared to type I RLN3-positive and type II NI neurons, and the MS-associated decrease. Statistical significance was determined using two-way ANOVA with a post hoc Tukey test: * MS effect, + neuronal type effect, # interaction of MS and neuronal type. The number of each symbol indicates the level of statistical significance: * ($p<0.05$), ** ($p<0.01$), *** ($p<0.001$). Please see **Supplementary Table 10** for details.

**Supplementary Figure 5. Number of counted cells in control and MS rats in the RNAscope studies**. **(a)** Total counted cells in the entire NI and its subregions, NIc and NId. **(b)** The percentage of counted cells expressing RLN3 mRNA in NI and its subregions. Statistical significance was assessed using a t-test, and revealed no sign
