## Supplementary Figures for "Early-life adversity alters adult nucleus incertus neurons: implications for neuronal mechanisms of increased stress and compulsive behavior vulnerability"

**Supplementary Figure 1.**

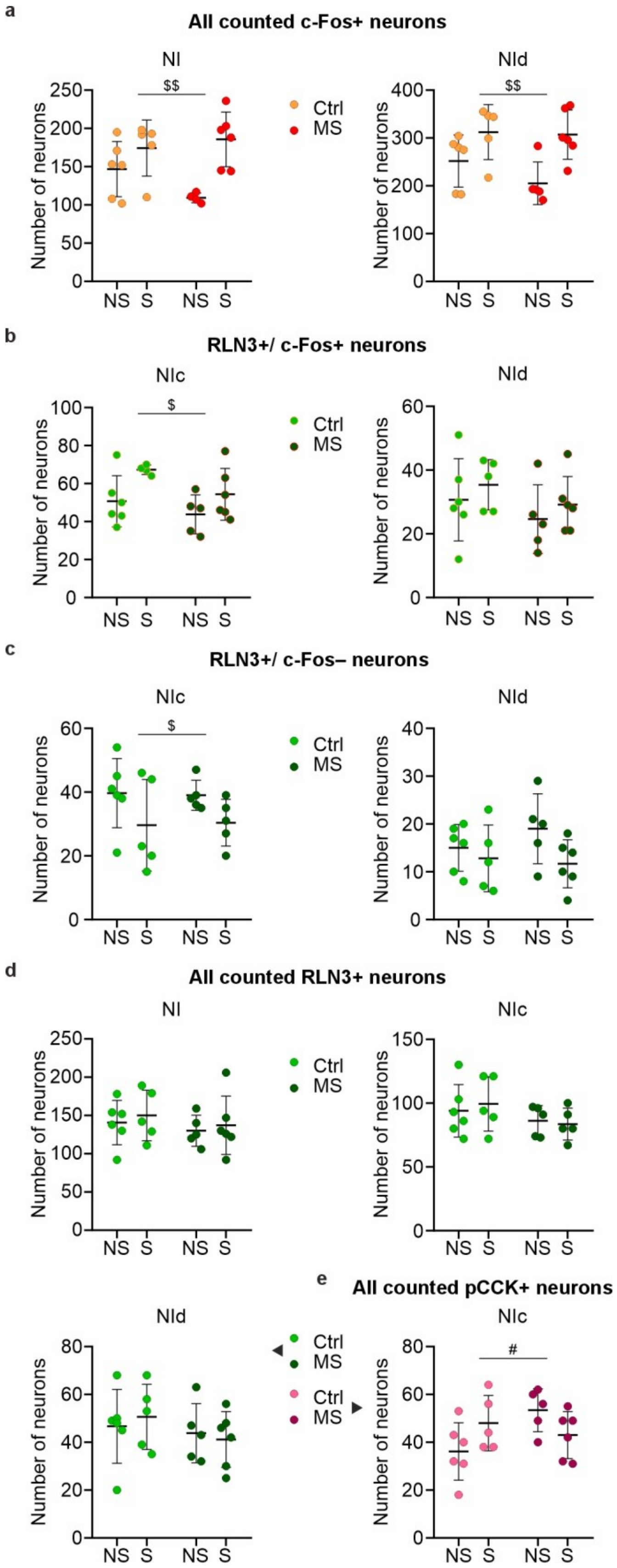

**Supplementary Figure 2.**

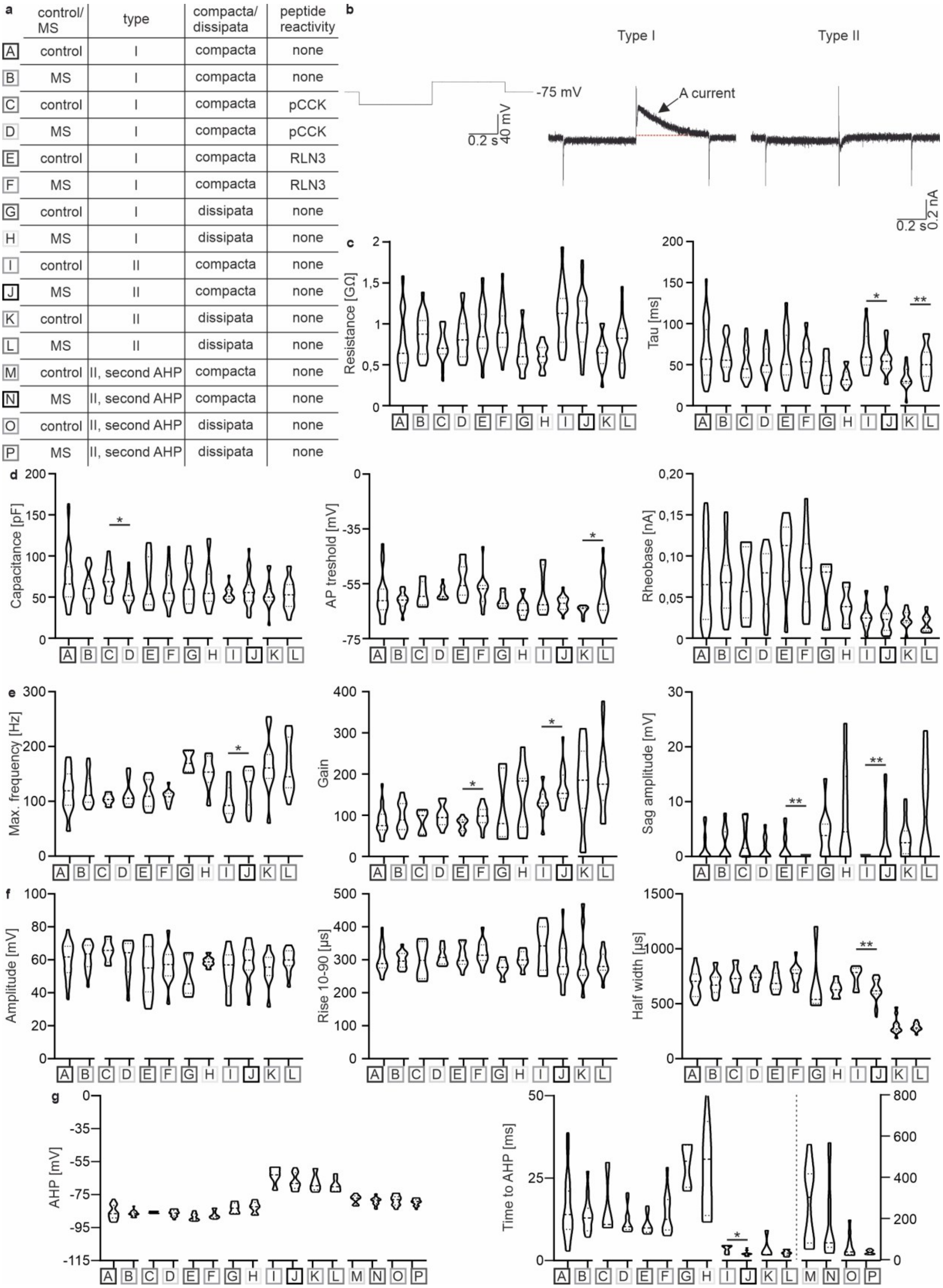


| | control/ MS | type | compacta/ dissipata | peptide reactivity |
|---|---|---|---|---|
| A | control | I | compacta | none |
| B | MS | I | compacta | none |
| C | control | I | compacta | pCCK |
| D | MS | I | compacta | pCCK |
| E | control | I | compacta | RLN3 |
| F | MS | I | compacta | RLN3 |
| G | control | I | dissipata | none |
| H | MS | I | dissipata | none |
| I | control | II | compacta | none |
| J | MS | II | compacta | none |
| K | control | II | dissipata | none |
| L | MS | II | dissipata | none |
| M | control | II, second AHP | compacta | none |
| N | MS | II, second AHP | compacta | none |
| O | control | II, second AHP | dissipata | none |
| P | MS | II, second AHP | dissipata | none |

**Supplementary Figure 3.**

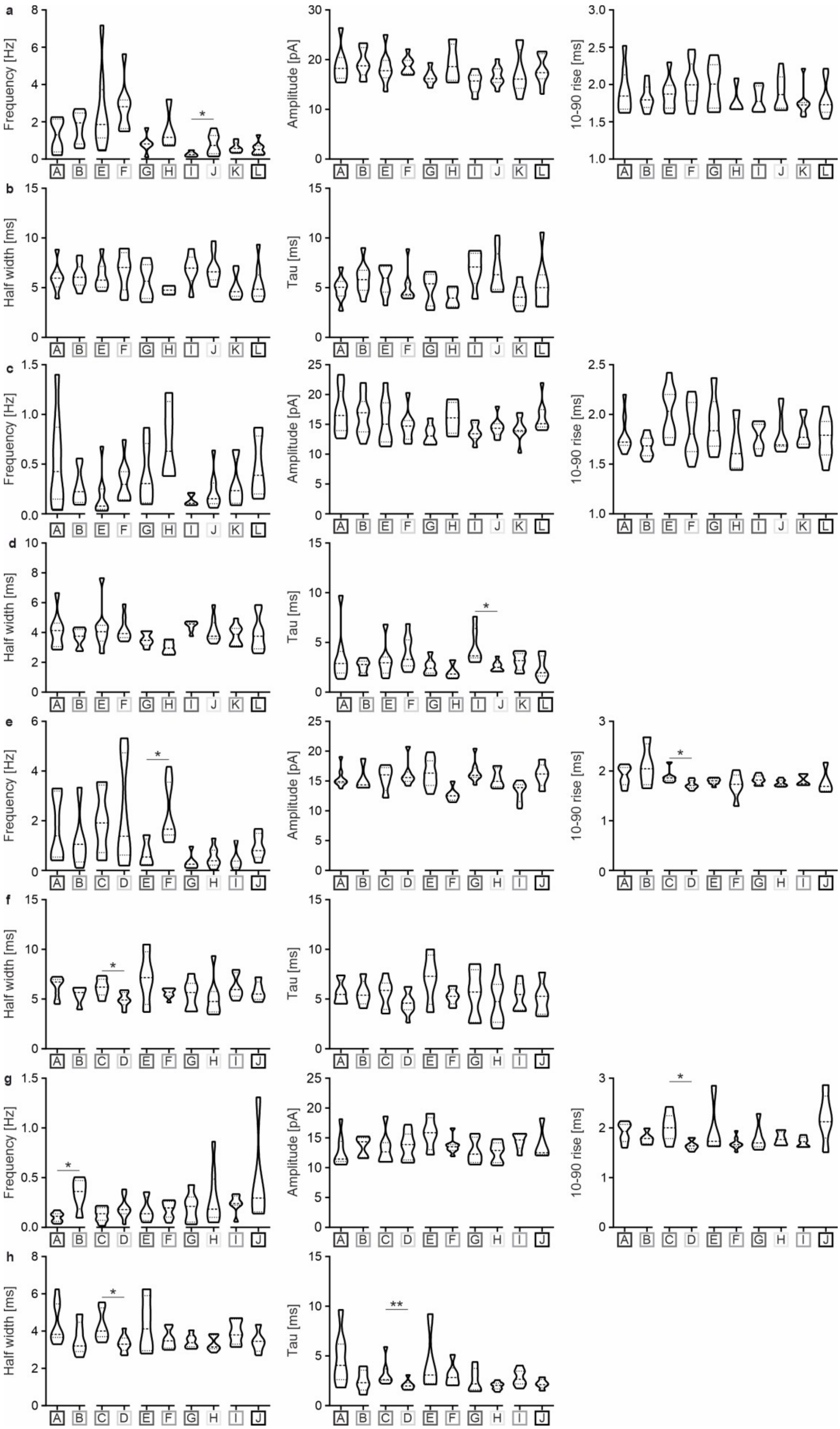

**Supplementary Figure 4.**

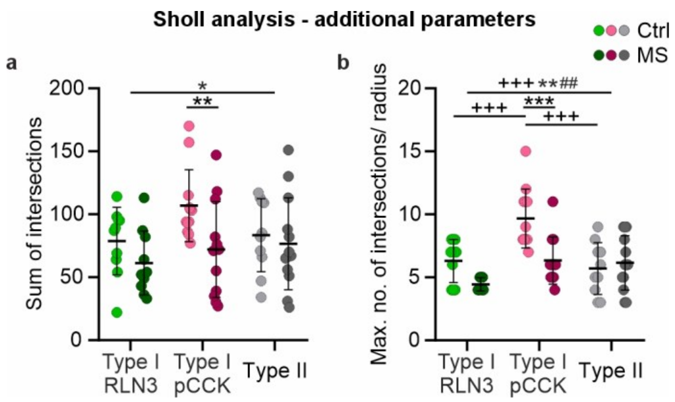


**Supplementary Figure 5.**

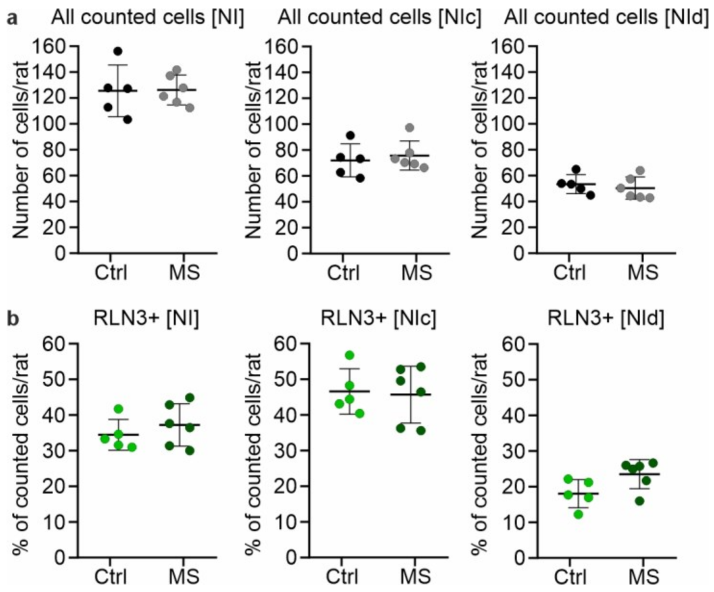
