## Supplementary Tables for "Early-life adversity alters adult nucleus incertus neurons: implications for neuronal mechanisms of increased stress and compulsive behavior vulnerability"

**Supplementary Table 1.** c-Fos expression in NI

| Location | Mean ± SD (n) | | | | Two-way ANOVA - effects (p and F values with df) | | | post hoc Tukey test (p, q and df values) | | | |
|---|---|---|---|---|---|---|---|---|---|---|---|
| | Ctrl | | MS | | Maternal separation | Restraint stress | Interaction | Ctrl NS vs. S NS | Ctrl NS vs. MS NS | MS NS vs. MS S | Ctrl S vs. MS S |
| | NS (6) | S (5) | NS (5) | S (6) | | | | | | | |
| All c-Fos+ cells | | | | | | | | | | | |
| NI | 252 ± 55 | 312 ± 58 | 205 ± 44 | 307 ± 51 | $F_{(1,18)}$ = 1.34, p = 0.26 | $F_{(1,18)}$ = 13.11, **p = 0.002** | $F_{(1,18)}$ = 1.86, p = 0.37 | - | - | - | - |
| NIc | 105 ± 24 | 138 ± 23 | 78 ± 7 | 121 ± 29 | $F_{(1,18)}$ = 4.81, **p = 0.04** | $F_{(1,18)}$ = 14.82, **p = 0.001** | $F_{(1,18)}$ = 0.25, p = 0.62 | - | - | - | - |
| NId | 147 ± 36 | 174 ± 37 | 109 ± 6 (4) | 186 ± 36 | $F_{(1,17)}$ = 0.81, p = 0.38 | $F_{(1, 17)}$ = 12.73, **p = 0.002** | $F_{(1,17)}$ = 2.84, p = 0.11 | - | - | - | - |
| RLN3+/cFos+ neurons | | | | | | | | | | | |
| NI | 81 ± 19 | 99 ± 15 | 69 ± 18 (4) | 84 ± 18 | $F_{(1,17)}$ = 6.21, **p = 0.02** | $F_{(1,17)}$ = 7.81, **p = 0.01** | $F_{(1,17)}$ = 0.15, p = 0.71 | - | - | - | - |
| NIc | 51 ± 13 | 63 ± 9 (4) | 44 ± 10 | 54 ± 14 | $F_{(1,17)}$ = 3.76, p = 0.07 | $F_{(1, 17)}$ = 7.05, **p = 0.02** | $F_{(1,17)}$ = 0.35, p = 0.56 | - | - | - | - |
| NId | 31 ± 13 | 35 ± 8 | 25 ± 11 | 29 ± 9 | $F_{(1,18)}$ = 1.92, p = 0.18 | $F_{(1,18)}$ = 1.10, p = 0.31 | $F_{(1,18)}$ = 0.0004, p = 0.99 | - | - | - | - |
| RLN3+/cFos- neurons | | | | | | | | | | | |
| NI | 55 ± 13 (5) | 42 ± 21 | 58 ± 11 | 49 ± 23 (5) | $F_{(1,16)}$ = 0.84, p = 0.78 | $F_{(1,16)}$ = 8.66, **p = 0.01** | $F_{(1,16)}$ = 0.0003, p = 0.99 | - | - | - | - |
| NIc | 40 ± 11 | 30 ± 14 | 39 ± 5 | 38 ± 19 (5) | $F_{(1,17)}$ = 0.0002, p = 0.99 | $F_{(1,17)}$ = 4.50, **p = 0.049** | $F_{(1,17)}$ = 0.03, p = 0.87 | - | - | - | - |

| | | | | | | | | | | | |
|---|---|---|---|---|---|---|---|---|---|---|---|
| NId | 15 ± 5 | 13 ± 7 | 19 ± 7 | 12 ± 5 | $F_{(1,18)}$ = 0.31, p = 0.59 | $F_{(1,18)}$ = 3.41, p = 0.08 | $F_{(1,18)}$ = 0.99, p = 0.33 | - | - | - | - |
| pCCK+/cFos+ neurons | | | | | | | | | | | |
| NIc | 4 ± 2 | 15 ± 9 | 4 ± 4 | 20 ± 9 | $F_{(1,18)}$ = 0.90, p = 0.36 | $F_{(1,18)}$ = 20.85, **p = 0.0002** | $F_{(1,18)}$ = 0.49, p = 0.49 | - | - | - | - |
| pCCK+/cFos- neurons | | | | | | | | | | | |
| NIc | 29 ± 10 | 26 ± 11 | 46 ± 8 | 19 ± 9 | $F_{(1,18)}$ = 1.39, p = 0.25 | $F_{(1,18)}$ = 12.72, **p = 0.002** | $F_{(1,18)}$ = 8.79, **p = 0.008** | $q_{(18)}$ = 0.60, p = 0.97 | $q_{(18)}$ = 0.60, **p = 0.04** | $q_{(18)}$ = 6.53, **p = 0.001** | $q_{(18)}$ = 1.79, p = 0.60 |

The symbol (-) indicates no post hoc analysis

**Supplementary Table 2**. Comparison of the proportions of c-Fos+ and c-Fos- neurons

| Number of c-Fos+ neurons/ total number of counted neurons | | | | | | | | Fisher's exact test (p value) | | | |
|---|---|---|---|---|---|---|---|---|---|---|---|
| RLN3 (NI) | | | | pCCK (NIc) | | | | RLN3: Ctrl NS vs. S | RLN3: MS NS vs. S | pCCK: Ctrl NS vs. S | pCCK: MS NS vs. S |
| Ctrl NS | Ctrl S | MS NS | MS S | Ctrl NS | Ctrl S | MS NS | MS S | | | | |
| 81/136 | 99/141 | 69/127 | 84/133 | 4/33 | 15/41 | 4/50 | 20/39 | 0.08 | 0.17 | **0.03** | **<0.0001** |

**Supplementary Table 3.** Active, passive membrane properties and spike properties – comparison of CCK+ and RLN3+ neurons

| Parameter | Mean/Median ± SD/iqr | | test (p and t/U values with df) | | |
|---|---|---|---|---|---|
| | C | | result | test | Welch's correction |
| | CCK (21) | RLN3 (15) | | | |
| resistance | 737,4 ± 187,2 | 911,8 ± 310,1 | $t_{(21.23)}$ = 1.94, p = 0.07 | t-test | yes |
| tau | 50,81 ± 20,18 | 59,88 ± 30,66 | $t_{(34)}$ = 1,072, p = 0.29 | t-test | no |
| capacitance | 68,68 ± 32,34 | 53,88 ± 58,39 | U = 140, p = 0.59 | M-W | no |
| AP threshold | -43,68 ± 4,56 | -38,66 ± 6,33 | $t_{(14)}$ = 1.58, p = 0.14 | t-test | no |
| rheobase | 6,57 ± 4,43 | 10,22 ± 4,27 | $t_{(14)}$ = 1.57, p = 0.14 | t-test | no |
| max FQ | 103,6 ± 10,46 | 113,1 ± 25,93 | $t_{(14)}$ = 0.78, p = 0.45 | t-test | no |
| gain | 85,47 ± 29,21 | 77,70 ± 15,88 | $t_{(14)}$ = 0.70, p = 0.50 | t-test | no |
| sag amplitude | 1,52 ± 6,23 | 0 ± 3,35 (13) | U = 105, p = 0.23 | M-W | no |
| amplitude | 65,50 ± 7,36 | 53,37 ± 14,96 | $t_{(13)}$ = 1.53, p = 0.15 | t-test | no |
| rise 10-90 | 298,9 ± 58,78 | 310,8 ± 35,59 | $t_{(13)}$ = 0.49, p = 0.64 | t-test | no |
| half width | 738,9 ± 121,8 | 701,3 ± 95,16 (14) | $t_{(13)}$ = 0.62, p = 0.55 | t-test | no |
| AHP | -70,80 ± 0,67 | -73,60 ± 2,174 | $t_{(13)}$ = 2.48, **p = 0.0278** | t-test | no |
| time to AHP | 11,02 ± 15,22 | 9,97 ± 4,47 (13) | U = 8, p = 0.15 | M-W | no |

**Supplementary Table 4.** Spontaneous and miniature postsynaptic currents properties – comparison of CCK+ and RLN3+ neurons

| Parameter | Mean/Median ± SD/iqr | | test (p and t/U values with df) | | |
|---|---|---|---|---|---|
| | C | | result | test | Welch's correction |
| | CCK n=7 | RLN3 n=4 | | | |
| mIPSC fq | 2,00 ± 1,22 | 0,68 ± 0,55 | $t_{(9)}$ = 2.01, p = 0.08 | t-test | no |
| mIPSC amplitude | 15,56 ± 6,56 | 14,34 ± 4,88 | U = 7, p = 0.23 | M-W | no |
| mIPSP 10-90 rise | 1,86 ± 0,42 | 2,05 ± 1,03 | U = 9, p = 0.41 | M-W | no |
| mIPSP half width | 6,14 ± 0,89 | 7,13 ± 2,80 | $t_{(3.353)}$ = 0.68, p = 0.54 | t-test | yes |
| mIPSP tau | 5,59 ± 1,44 | 7,07 ± 2,62 | $t_{(9)}$ = 1.23, p = 0.25 | t-test | no |
| mEPSP fq | 0,13 ± 0,07 | 0,17 ± 0,13 | $t_{(9)}$ = 0.73, p = 0.48 | t-test | no |
| mEPSP amplitude | 13,22 ± 2,74 | 15,76 ± 2,83 | $t_{(9)}$ = 1.46, p = 0.18 | t-test | no |
| mEPSP 10-90 rise | 2,00 ± 0,81 | 1,73 ± 1,22 | U = 10, p = 0.53 | M-W | no |
| mEPSP half width | 4,27 ± 0,83 | 4,33 ± 1,55 | $t_{(9)}$ = 0.08, p = 0.94 | t-test | no |
| mEPSP tau | 3,22 ± 1,28 | 4,38 ± 3,30 | U = 14, p > 0.99 | t-test | no |

**Supplementary Table 5**. Active, passive membrane properties and spike properties – two-way ANOVA analysis

| Parameter | Mean ± SD | | | | Two-way ANOVA - effects (p and F values with df) | | | | post hoc Tukey test (p, t and df values) | | | |
|---|---|---|---|---|---|---|---|---|---|---|---|---|
| | C | | MS | | Maternal separation | Type | Interaction | White adjustment | I C vs. I MS | I C vs. II C | II C vs. II MS | I MS vs. II MS |
| | Type I (n) | Type II (n) | Type I (n) | Type II (n) | | | | | | | | |
| resistance | 751,9 ± 287,2 (89) | 868,0 ± 374,1 (52) | 794,4 ± 272,3 (105) | 887,9 ± 336,9 (59) | $F_{(1,301)}$ = 0.72, p = 0.40 | $F_{(1,301)}$ = 8.10, **p = 0.0047** | $F_{(1,301)}$ = 0.09, p = 0.76 | no | - | - | - | - |
| tau | 52,17 ± 25,30 (89) | 45,50 ± 21,16 (52) | 45,61 ± 19,53 (105) | 48,22 ± 24,03 (59) | $F_{(1,296)}$ = 0.50, p = 0.48 | $F_{(1,296)}$ = 0.56, p = 0.46 | $F_{(1,301)}$ = 2.92, p = 0.09 | no | - | - | - | - |
| capacitance | 70,95 ± 27,89 (89) | 52,49 ± 14,62 (52) | 59,67 ± 20,80 (105) | 54,58 ± 18,69 (59) | $F_{(1,296)}$ = 3.00, p = 0.08 | $F_{(1,296)}$ = 19.66, **p < 0.0001** | $F_{(1,296)}$ = 6.35, **p = 0.0123** | no | $t_{(296)}$ = 3.55, **p = 0.0027** | $t_{(296)}$ = 4.72, **p < 0.0001** | $t_{(296)}$ = 0.49, p = 0.99 | $t_{(296)}$ = 1.42, p = 0.64 |
| AP threshold | -42,89 ± 6,88 (45) | -45,44 ± 7,337 (30) | -43,99 ± 6,58 (65) | -45,27 ± 6,19 (46) | $F_{(1,176)}$ = 2.82, p = 0.10 | $F_{(1,176)}$ = 4.88, **p = 0.0285** | $F_{(1,176)}$ = 0.40, p = 0.53 | no | - | - | - | - |
| rheobase | 75,80 ± 47,47 (42) | 24,47 ± 12,39 (23) | 69,03 ± 39,70 (53) | 20,51 ± 14,12 (41) | $F_{(1,301)}$ = 0.87, p = 0.35 | $F_{(1,301)}$ = 75.3, **p < 0.0001** | $F_{(1,301)}$ = 0.06, p = 0.81 | no | - | - | - | - |
| max FQ | 120,9 ± 34,90 (45) | 128,6 ± 51,33 (30) | 123,3 ± 33,02 (65) | 142,9 ± 43,60 (46) | $F_{(1,301)}$ = 13.10, p = 0.32 | $F_{(1,301)}$ = 0.01, p = 0.16 | $F_{(1,301)}$ = 0.42, p = 0.27 | yes | - | - | - | - |
| gain | 90,3 ± 43,48 (43) | 153,2 ± 75,91 (53) | 103,4 ± 42,15 (23) | 176,9 ± 61,72 (40) | $F_{(1,155)}$ = 3.60, p = 0.11 | $F_{(1,155)}$ = 79.40, **p < 0.0001** | $F_{(1,155)}$ = 0.44, p = 0.69 | yes | - | - | - | - |
| sag amplitude | 6,00 ± 3,71 (44) | 4,30 ± 2,92 (17) | 6,18 ± 3,64 (34) | 11,87 ± 7,56 (27) | $F_{(1,115)}$ = 17.24, **p < 0.0001** | $F_{(1,115)}$ = 4.56, **p = 0.0349** | $F_{(1,115)}$ = 15.64, **p = 0.0001** | no | $t_{(115)}$ = 0.23, p = 0.99 | $t_{(115)}$ = 1.73, p = 0.61 | $t_{(115)}$ = 7.12, **p < 0.0001** | $t_{(115)}$ = 6.46, **p < 0.0001** |
| amplitude | 57,54 ± 12,48 (40) | 54,27 ± 10,80 (52) | 59,41 ± 9,26 (20) | 58,63 ± 9,33 (37) | $F_{(1,144)}$ = 2.92, p = 0.09 | $F_{(1,144)}$ = 1.23, p = 0.27 | $F_{(1,144)}$ = 0.47, p = 0.50 | no | - | - | - | - |
| rise 10-90 | 300,1 ± 39,80 (40) | 315,2 ± 76,12 (52) | 308,5 ± 28,80 (21) | 289,0 ± 51,99 (37) | $F_{(1,143)}$ = 1.21, p = 0.41 | $F_{(1,143)}$ = 0.07, p = 0.84 | $F_{(1,143)}$ = 4.53, p = 0.11 | yes | - | - | - | - |

| | | | | | | | | | | | | |
|---|---|---|---|---|---|---|---|---|---|---|---|---|
| half width | 679,3 ± 116,2 (40) | 486,5 ± 242,3 (52) | 710,4 ± 95,41 (21) | 508,9 ± 209,2 (37) | $F_{(1,141)}$ = 0.89, p = 0.45 | $F_{(1,141)}$ = 48.70, **p < 0.0001** | $F_{(1,141)}$ = 0.02, p = 0.90 | yes | - | - | - | - |
| AHP | -71,28 ± 3,730 (40) | -63,16 ± 2,85 (52) | -71,14 ± 2,60 (21) | -64,28 ± 2,23 (36) | $F_{(1,146)}$ = 0.95, p = 0.33 | $F_{(1,146)}$ = 224.30, **p < 0.0001** | $F_{(1,146)}$ = 1.57, p = 0.21 | no | - | - | - | - |
| time to AHP | 17,41 ± 10,69 (40) | 152,9 ± 151,4 (52) | 13,96 ± 5,61 (21) | 72,65 ± 54,15 (37) | $F_{(1,128)}$ = 0.06, p = 0.08 | $F_{(1,128)}$ = 60.00, **p < 0.0001** | $F_{(1,128)}$ = 11.61, **p = 0.0008** | yes | $t_{(128)}$ = 0.25, p = 0.99 | $t_{(128)}$ = -7.75, **p < 0.0001** | $t_{(128)}$ = 4.35, **p = 0.0002** | $t_{(128)}$ = -3.94, **p = 0.0008** |

The symbol (-) indicates no post hoc analysis

**Supplementary Table 6.** Sag occurrence in CCK+ and RLN3+ neurons

| noumber of sag+ neurons/total noumber of neurons | | | | | | Fisher's exact test (p value) | | | | |
|---|---|---|---|---|---|---|---|---|---|---|
| C | | C | | MS | | CCK vs. RLN3 | I C vs. I MS | I C vs. II C | I MS vs. II MS | II MS vs. II C |
| CCK | RLN3 | I | II | I | II | | | | | |
| 11/21 | 6/15 | 46/89 | 22/46 | 38/92 | 36/59 | 0.52 | 0.18 | 0.72 | 0.24 | **0.0203** |

**Supplementary Table 7**. Active, passive membrane properties and spike properties – t-test

| Parameter | Mean/Median ± SD/iqr | | | | | | | | | | | | | | | | t-test (p and t/U values with df) | | | | | | | |
|---|---|---|---|---|---|---|---|---|---|---|---|---|---|---|---|---|---|---|---|---|---|---|---|---|
| | group | | | | | | | | | | | | | | | | | | | | | | | |
| | A (36) | B (23) | C (21) | D (22) | E (15) | F (27) | G (12) | H (16) | I (19) | J (30) | K (17) | L (20) | M | N | O | P | A vs. B | C vs. D | E vs. F | G vs. H | I vs. J | K vs. L | M vs. N | O vs. P |
| resistance | 639,6 ± 398,8 | 874,1 ± 410,7 | 737,4 ± 187,2 | 835,5 ± 247,1 | 911,8 ± 310,1 | 915,7 ± 268,4 | 642,4 ± 232,1 | 611,8 ± 144,4 (15) | 1109 ± 356,2 | 1016 ± 324,9 | 635 ± 193,9 | 792,4 ± 278,4 | - | - | - | - | U = 291, p = 0.06, M-W | $t_{(41)}$ = 1.46, p = 0.15, t-test | $t_{(40)}$ = 0.04, p = 0.97, t-test | $t_{(25)}$ = 0.42, p = 0.68, t-test | $t_{(47)}$ = 0.94, p = 0.35, t-test | $t_{(35)}$ = 1.96, p = 0.06, t-test | - | - |
| tau | 66,26 ± 33,14 | 60,09 ± 20,22 | 50,81 ± 20,18 | 53,29 ± 18,15 | 59,88 ± 30,66 | 56,17 ± 20,20 (26) | 39,90 ± 17,12 | 34,06 ± 10,73 (13) | 52,17 ± 25,30 | 54,95 ± 16,70 (29) | 33,96 ± 12,83 | 51,41 ± 20,61 | - | - | - | - | $t_{(56,92)}$ = 0.89, p = 0.38, Welch | $t_{(41)}$ = 0.42, p = 0.67, t-test | $t_{(39)}$ = 0.47, p = 0.64, t-test | $t_{(23)}$ = 1.03, p = 0.31, t-test | $t_{(46)}$ = 2.03, **p = 0.0484**, t-test | $t_{(35)}$ = 3.02, **p = 0.0046**, t-test | - | - |
| capacitance | 65,94 ± 36,60 | 60,49 ± 22,54 | 68,81 ± 18,33 | 55,88 ± 15,03 | 53,88 ± 58,39 | 54,82 ± 30,61 (26) | 64,62 ± 26,22 | 62,43 ± 26,74 | 51,16 ± 11,61 (16) | 55,59 ± 19,64 | 53,00 ± 16,66 | 52,68 ± 17,57 | - | - | - | - | U = 336, p = 0.23, M-W | $t_{(41)}$ = 2.54, **p = 0.0152**, t-test | U = 185, p = 0.80, M-W | $t_{(26)}$ = 0.22, p = 0.83, t-test | U = 233, p = 0.88, Mann-Whitney | $t_{(35)}$ = 0.06, p = 0.95, t-test | - | - |
| AP threshold | -46,16 ± 7,54 | 45,93 ± 3,8 (20) | -44,60 ± 8,57 | -44,66 ± 3,78 (21) | -40,67 ± 10,31 | -41,84 ± 5,85 | -45,89 ± 2,90 | -48,68 ± 3,66 | -47,68 ± 16,40 | -47,14 ± 4,17 (28) | -49,78 ± 1,81 (16) | -44,51 ± 7,40 | - | - | - | - | U = 162, p = 0.94, M-W | U = 28, p = 0.88, M-W | U = 60, p = 0.26, M-W | $t_{(10)}$ = 1.41, p = 0.19, t-test | U = 112, p = 0.75, M-W | $t_{(18.85)}$ = 2.81, **p = 0.0113**, Welch | - | - |
| rheobase | 69,15 ± 49,45 | 67,71 ± 39,55 | 65,67 ± 44,27 | 71,58 ± 35,18 | 102,20 ± 42,71 | 82,60 ± 45,16 | 80,41 ± 74,31 | 38,45 ± 45,01 | 24,57 ± 15,55 | 22,96 ± 20,34 | 21,21 ± 12,13 | 12,36 ± 16,25 | - | - | - | - | $t_{(37)}$ = 0.10, p = 0.92, t-test | $t_{(16)}$ = 0.2978, p = 0.7697, t-test | $t_{(24)}$ = 1.118, p = 0.2748, t-test | U = 13, p = 0.5303, Mann-Whitney | U = 117.5, p = 0.6177, Mann-Whitney | U = 70, p = 0.1620, Mann-Whitney | - | - |
| max FQ | 119,1 ± 56,83 | 110,8 ± 43,44 | 103,3 ± 18,65 | 105,6 ± 33,09 | 113,1 ± 25,93 | 107,5 ± 12,15 | 172,0 ± 18,84 | 151,2 ± 31,95 | 92,27 ± 47,19 | 136,1 ± 62,32 | 165,8 ± 50,66 | 163,6 ± 47,81 | - | - | - | - | U = 194, p = 0.93, M-W | U = 129, p = 0.63, M-W | $t_{(13.44)}$ = 0.66, p = 0.52, Welch | $t_{(10)}$ = 1.29, p = 0.23, t-test | U = 74, **p = 0.0397**, M-W | $t_{(27)}$ = 0.12, p = 0.90, t-test | - | - |
| gain | 85,00 ± 33,14 (35) | 95,92 ± 35,45 | 85,47 ± 29,21 | 98,96 ± 26,14 | 77,70 ± 15,88 | 96,87 ± 25,23 | 121,10 ± 86,57 | 144,70 ± 81,69 | 129,70 ± 28,20 | 153,20 ± 56,50 | 174,60 ± 96,85 | 189,60 ± 81,26 | - | - | - | - | $t_{(37)}$ = 0.99, p = 0.33, t-test | $t_{(16)}$ = 0.95, p = 0.36, t-test | $t_{(24)}$ = 2.21, **p = 0.0367**, t-test | $t_{(10)}$ = 0.48, p = 0.64, t-test | U = 60.50, **p = 0.0138**, M-W | $t_{(27)}$ = 0.45, p = 0.65, t-test | - | - |
| sag amplitude | 0 ± 2,92 (34) | 0 ± 4,51 (21) | 1,52 ± 6,23 | 0 ± 3,32 (21) | 0 ± 3,36 (13) | 0 ± 0 (21) | 3,84 ± 5,92 | 4,52 ± 14,62 | 0 ± 0 (15) | 0 ± 6,45 (28) | 2,51 ± 4,15 | 7,20 ± 15,91 | - | - | - | - | U = 183, p = 0.44, M-W | U = 179, p = 0.26, M-W | U = 94.50, **p = 0.0154**, M-W | U = 80, p = 0.46, M-W | U = 112.5, **p = 0.0054**, M-W | U = 133, p = 0.26, M-W | - | - |
| | **A (20)** | **B (16)** | **C (4)** | **D (13)** | **E (11)** | **F (15)** | **G (5)** | **H (7)** | **I (10)** | **J (22)** | **K (11)** | **L (15)** | **M (5)** | **N (11)** | **O (8)** | **P (15)** | | | | | | | | |
| amplitude | 60,08 ± 10,86 | 61,71 ± 8,35 (15) | 65,50 ± 7,36 | 60,60 ± 11,43 | 53,37 ± 14,96 | 56,29 ± 10,14 | 50,18 ± 12,09 | 58,95 ± 3,26 | 53,44 ± 12,35 (9) | 58,14 ± 10,68 | 54,94 ± 9,93 | 59,39 ± 7,00 (14) | - | - | - | - | $t_{(34)}$ = 0.50, p = 0.62, t-test | $t_{(15)}$ = 0.80, p = 0.44, t-test | $t_{(24)}$ = 0.59, p = 0.56, t-test | $t_{(4.42)}$ = 1.58, p = 0.18, Welch | $t_{(29)}$ = 1.07, p = 0.306, t-test | $t_{(23)}$ = 1.31, p = 0.20, t-test | - | - |
| rise 10-90 | 300,6 ± 40,56 | 299,4 ± 24,00 | 298,9 ± 58,78 | 309,9 ± 26,01 | 310,8 ± 35,59 | 321,0 ± 34,31 | 275,4 ± 28,10 | 301,1 ± 26,92 | 338,6 ± 66,77 (8) | 290,7 ± 60,67 | 298,2 ± 80,96 | 286,2 ± 36,30 (14) | - | - | - | - | $t_{(31.52)}$ = 0.11, p = 0.913, Welch | $t_{(3.369)}$ = 0.36, p = 0.74, Welch | $t_{(24)}$ = 0.73, p = 0.47, t-test | $t_{(10)}$ = 1.60, p = 0.14, t-test | $t_{(28)}$ = 1.86, p = 0.07, t-test | $t_{(13.16)}$ = 0.45, p = 0.65, Welch | - | - |
| half width | 683,2 ± 119,10 | 688,7 ± 99,86 | 738,9 ± 121,80 | 727,1 ± 67,69 | 701,3 ± 95,16 (10) | 758,9 ± 97,61 (14) | 539,8 ± 416,80 | 626,7 ± 200,50 | 784,9 ± 198,50 (9) | 618,4 ± 64,70 | 612,4 ± 279,40 | 601,9 ± 69,00 | - | - | - | - | $t_{(35)}$ = 0.15, p = 0.88, t-test | $t_{(15)}$ = 0.25, p = 0.80, t-test | $t_{(22)}$ = 1.44, p = 0.16, t-test | U = 11, p = 0.34, M-W | U = 32, **p = 0.0072**, M-W | U = 76, p = 0.76, M-W | - | - |
| AHP | -70,87 ± 4,24 | -71,49 ± 1,73 | -70,80 ± 0,67 | -71,29 ± 1,95 | -74,62 ± 3,75 | -72,08 ± 4,10 | -68,23 ± 3,23 | -67,73 ± 3,48 | -48,27 ± 5,70 | -62,84 ± 2,66 | -52,19 ± 4,91 | -64,07 ± 2,37 | -52,81 ± 5,54 | -63,46 ± 3,12 | -54,50 ± 3,94 | -64,58 ± 2,06 | $t_{(26.01)}$ = 0.60, p = 0.56, Welch | $t_{(15)}$ = 0.48, p = 0.64, t-test | U = 55, p = 0.16, M-W | $t_{(10)}$ = 0.25, p = 0.81, t-test | $t_{(16)}$ = 1.46, p = 0.16, t-test | $t_{(30)}$ = 1.31, p = 0.20, t-test | $t_{(11)}$ = 0.65, p = 0.53, t-test | $t_{(24)}$ = 1.11, p = 0.28, t-test |
| | **A (20)** | **B (16)** | **C (4)** | **D (13)** | **E (11)** | **F (15)** | **G (5)** | **H (7)** | **I (10)** | **J (22)** | **K (11)** | **L (15)** | **M (10)** | **N (22)** | **O (11)** | **P (15)** | | | | | | | | |
| time to AHP | 15,81 ± 9,60 (19) | 13,95 ± 5,27 (14) | 11,02 ± 15,22 | 10,35 ± 3,97 (12) | 10,90 ± 2,94 (9) | 14,34 ± 6,10 (14) | 30,17 ± 12,76 | 30,79 ± 28,49 | 4,36 ± 2,26 | 2,07 ± 0,45 | 1,94 ± 4,39 | 2,37 ± 1,50 | 306,30 ± 335,97 | 85,61 ± 227,82 | 41,52 ± 73,58 (10) | 39,69 ± 19,16 (12) | $t_{(28.94)}$ = 0.72, p = 0.48, Welch | U = 16, p = 0.38, M-W | $t_{(19.89)}$ = 1.81, p = 0.09, Welch | U = 16, p = 0.88, M-W | U = 12, **p = 0.0460**, M-W | U = 20, p > 0.99, M-W | U = 74, p = 0.15, M-W | U = 58, p = 0.92, M-W |

**Supplementary Table 8**. Spontaneous and miniature postsynaptic currents properties – two-way ANOVA analysis

| Parameter | Mean ± SD | | | | Two-way ANOVA - effects (p and F values with df) | | | | post hoc Tukey test (p, t and df values) | | | |
|---|---|---|---|---|---|---|---|---|---|---|---|---|
| | C | | MS | | Maternal separation | Type | Interaction | White adjustment | I C vs. I MS | I C vs. II C | II C vs. II MS | I MS vs. II MS |
| | Type I (n) | Type II (n) | Type I (n) | Type II (n) | | | | | | | | |
| sIPSC fq | 1,35 ± 0,96 (21) | 0,45 ± 0,28 (14) | 2,10 ± 1,19 (20) | 0,78 ± 0,59 (18) | $F_{(1,69)}$ = 6.54, p = 0.11 | $F_{(1,69)}$ = 27.92, **p < 0.0001** | $F_{(1,69)}$ = 1.00, p = 0.52 | yes | - | - | - | - |
| sIPSP amplitude | 18,09 ± 3,08 (22) | 16,50 ± 3,54 (14) | 19,12 ± 2,58 (21) | 17,32 ± 2,49 (18) | $F_{(1,71)}$ = 1.84, p = 0.18 | $F_{(1,71)}$ = 6.18, **p = 0.0153** | $F_{(1,71)}$ = 0.03, p = 0.87 | no | - | - | - | - |
| sIPSP 10-90 rise | 1,92 ± 0,27 (22) | 1,78 ± 0,18 (14) | 1,89 ± 0,24 (21) | 1,84 ± 0,23 (18) | $F_{(1,71)}$ = 0.07, p = 0.79 | $F_{(1,71)}$ = 2.82, p = 0.10 | $F_{(1,71)}$ = 0.90, p = 0.34 | no | - | - | - | - |
| sIPSP half width | 5,96 ± 1,47 (22) | 5,81 ± 1,64 (14) | 6,08 ± 1,54 (21) | 6,13 ± 1,79 (18) | $F_{(1,71)}$ = 0.34, p = 0.56 | $F_{(1,71)}$ = 0.02, p = 0.90 | $F_{(1,71)}$ = 0.07, p = 0.79 | no | - | - | - | - |
| sIPSP tau | 5,21 ± 1,43 (22) | 5,30 ± 1,96 (14) | 5,23 ± 1,61 (21) | 5,92 ± 2,26 (18) | $F_{(1,71)}$ = 0.56, p = 0.46 | $F_{(1,71)}$ = 0.85, p = 0.36 | $F_{(1,71)}$ = 0.49, p = 0.49 | no | - | - | - | - |
| sEPSP fq | 0,32 ± 0,29 (21) | 0,19 ± 0,15 (13) | 0,33 ± 0,28 (20) | 0,31 ± 0,24 (17) | $F_{(1,60)}$ = 0.20, p = 0.66 | $F_{(1,60)}$ = 1.93, p = 0.17 | $F_{(1,60)}$ = 0.21, p = 0.65 | no | - | - | - | - |
| sEPSP amplitude | 15,50 ± 3,54 (22) | 14,01 ± 1,84 (14) | 15,70 ± 2,92 (21) | 15,36 ± 2,24 (18) | $F_{(1,71)}$ = 1.39, p = 0.24 | $F_{(1,71)}$ = 1.94, p = 0.17 | $F_{(1,71)}$ = 0.77, p = 0.38 | no | - | - | - | - |
| sEPSP 10-90 rise | 1,90 ± 0,26 (22) | 1,80 ± 0,14 (14) | 1,74 ± 0,23 (21) | 1,79 ± 0,20 (18) | $F_{(1,71)}$ = 2.96, p = 0.09 | $F_{(1,71)}$ = 0.36, p = 0.55 | $F_{(1,71)}$ = 2.23, p = 0.14 | no | - | - | - | - |
| sEPSP half width | 3,85 ± 0,89 (21) | 4,07 ± 0,64 (14) | 3,76 ± 0,78 (21) | 4,01 ± 0,98 (18) | $F_{(1,70)}$ = 0.15, p = 0.70 | $F_{(1,70)}$ = 1.42, p = 0.24 | $F_{(1,70)}$ = 0.01, p = 0.93 | no | - | - | - | - |

| | | | | | | | | | | | | |
|---|---|---|---|---|---|---|---|---|---|---|---|---|
| sEPSP tau | 2,60 ± 0,86 (19) | 3,45 ± 1,04 (14) | 3,00 ± 1,36 (20) | 2,50 ± 0,87 (16) | $F_{(1,65)}$ = 1.11, p = 0.30 | $F_{(1,65)}$ = 0.43, p = 0.51 | $F_{(1,65)}$ = 6,57, **p = 0.0127** | no | $t_{(65)}$ = 1.65, p = 0.65 | $t_{(65)}$ = 3.10, p = 0.14 | $t_{(65)}$ = 3.35, p = 0.09 | $t_{(65)}$ = 1.99, p = 0.50 |
| mIPSC fq | 1,30 ± 1,19 (25) | 0,65 ± 0,31 (7) | 1,29 ± 1,15 (30) | 0,97 ± 0,53 (5) | $F_{(1,60)}$ = 0.20, p = 0.66 | $F_{(1,60)}$ = 1.93, p = 0.17 | F(1,60) = 0.21, p = 0.65 | no | - | - | - | - |
| mIPSC amplitude | 14,97 ± 2,17 (25) | 15,39 ± 1,54 (7) | 15,36 ± 2,34 (30) | 16,02 ± 1,97 (5) | $F_{(1,63)}$ = 0.52, p = 0.47 | $F_{(1,63)}$ = 0.59, p = 0.45 | $F_{(1,63)}$ = 0.03, p = 0.86 | no | - | - | - | - |
| mIPSP 10-90 rise | 1,87 ± 0,26 (25) | 1,76 ± 0,07 (7) | 1,78 ± 0,12 (30) | 1,81 ± 0,11 (5) | $F_{(1,63)}$ = 0.22, p = 0.64 | $F_{(1,63)}$ = 0.54, p = 0.46 | $F_{(1,63)}$ = 1.79, p = 0.19 | no | - | - | - | - |
| mIPSP half width | 6,15 ± 1,51 (25) | 6,17 ± 1,18 (7) | 5,10 ± 0,84 (30) | 5,78 ± 0,97 (5) | $F_{(1,62)}$ = 3.54, p = 0.06 | $F_{(1,62)}$ = 0.85, p = 0.36 | $F_{(1,62)}$ = 0.73, p = 0.40 | no | - | - | - | - |
| mIPSP tau | 5,83 ± 1,83 (25) | 5,25 ± 1,41 (6) | 4,99 ± 1,47 (30) | 5,05 ± 1,74 (5) | $F_{(1,63)}$ = 0.97, p = 0.33 | $F_{(1,63)}$ = 0.24, p = 0.63 | $F_{(1,63)}$ = 0.37, p = 0.55 | no | - | - | - | - |
| sEPSP fq | 0,16 ± 0,10 (24) | 0,24 ± 0,09 (6) | 0,22 ± 0,13 (30) | 0,47 ± 0,48 (5) | $F_{(1,61)}$ = 7.82, **p = 0.0069** | $F_{(1,61)}$ = 9.93, **p = 0.0025** | $F_{(1,61)}$ = 2.06, p = 0.16 | no | - | - | - | - |
| sEPSP amplitude | 13,36 ± 2,64 (25) | 14,43 ± 1,37 (7) | 13,37 ± 1,78 (30) | 13,89 ± 2,61 (5) | $F_{(1,63)}$ = 0.15, p = 0.70 | $F_{(1,63)}$ = 1.28, p = 0.26 | $F_{(1,63)}$ = 0.15, p = 0.70 | no | - | - | - | - |
| sEPSP 10-90 rise | 1,87 ± 0,28 (23) | 1,72 ± 0,10 (7) | 1,72 ± 0,12 (30) | 1,65 ± 0,11 (5) | $F_{(1,61)}$ = 2.95, p = 0.09 | $F_{(1,61)}$ = 3.08, p = 0.08 | $F_{(1,61)}$ = 0.33, p = 0.57 | no | - | - | - | - |
| sEPSP half width | 4,13 ± 1,00 (25) | 3,95 ± 0,61 (7) | 3,43 ± 0,53 (30) | 3,43 ± 0,61 (5) | $F_{(1,63)}$ = 6.24, **p = 0.0151** | $F_{(1,63)}$ = 0.14, p = 0.71 | $F_{(1,63)}$ = 0.13, p = 0.72 | no | - | - | - | - |
| sEPSP tau | 3,18 ± 1,39 (22) | 2,82 ± 0,84 (5) | 2,43 ± 0,94 (30) | 2,20 ± 0,50 (5) | $F_{(1,58)}$ = 3.74, p = 0.06 | $F_{(1,58)}$ = 0.69, p = 0.41 | $F_{(1,58)}$ = 0.04, p = 0.84 | no | - | - | - | - |

The symbol (-) indicates no post hoc analysis

**Supplementary Table 9.** Spontaneous and miniature postsynaptic currents properties – t-test

| Parameter | Mean/Median ± SD/iqr | | | | | | | | | | | | t-test (p and t/U values with df) | | | | | |
|---|---|---|---|---|---|---|---|---|---|---|---|---|---|---|---|---|---|---|
| | group | | | | | | | | | | | | A vs. B | C vs. D | E vs. F | G vs. H | I vs. J | K vs. L |
| | A (8) | B (9) | C | D | E (8) | F(8) | G(6) | H (4) | I (6) | J (9) | K (8) | L (9) | | | | | | |
| sIPSC fq | 1,28 ± 0,86 | 1,73 ± 0,83 | - | - | 2,55 ± 2,17 | 2,80 ± 1,35 | 0,83 ± 0,51 | 1,56 ± 1,12 | 0,27 ± 0,13 | 0,80 ± 0,55 | 0,61 ± 0,27 (7) | 0,56 ± 0,35 (8) | $t_{(15)}$ = 1.10, p = 0.29, t-test | - | $t_{(14)}$ = 0.27, p = 0.79, t-test | $t_{(8)}$ = 1.43, p = 0.19, t-test | $t_{(9.333)}$ = 2.82, **p = 0.0194**, Welch | $t_{(13)}$ = 0.30, p = 0.77, t-test |
| sIPSC amplitude | 18,94 ± 3,49 | 19,36 ± 2,86 | - | - | 18,39 ± 3,35 | 18,83 ± 1,79 | 16,56 ± 1,70 | 19,19 ± 3,82 | 15,36 ± 2,15 | 16,68 ± 1,97 | 17,36 ± 4,25 | 17,95 ± 2,91 | $t_{(15)}$ = 0.27, p = 0.79, t-test | - | $t_{(14)}$ = 0.33, p = 0.75, t-test | $t_{(8)}$ = 1.51, p = 0.17, t-test | $t_{(13)}$ = 1.23, p = 0.24, t-test | $t_{(15)}$ = 0.34, p = 0.74, t-test |
| sIPSP 10-90 rise | 1,92 ± 0,31 | 1,83 ± 0,16 | - | - | 1,87 ± 0,22 | 2,01 ± 0,29 | 2,01 ± 0,58 | 1,68 ± 0,35 | 1,80 ± 0,17 | 1,89 ± 0,23 | 1,76 ± 0,12 | 1,80 ± 0,35 | $t_{(15)}$ = 0.79, p = 0.44, t-test | - | $t_{(14)}$ = 1.03, p = 0.32, t-test | U = 6, p = 0.23, M-W | $t_{(13)}$ = 0.83, p = 0.42, t-test | U = 34, p = 0.87, M-W |
| sIPSP half width | 5,98 ± 1,44 | 6,14 ± 1,21 | - | - | 6,16 ± 1,42 | 6,69 ± 1,90 | 5,67 ± 1,80 | 4,75 ± 0,39 | 6,83 ± 1,63 | 6,91 ± 1,49 | 5,06 ± 1,25 | 5,36 ± 1,80 | $t_{(15)}$ = 0.25, p = 0.81, t-test | - | $t_{(14)}$ = 0.63, p = 0.54, t-test | $t_{(5,672)}$ = 1.21, p = 0.28, Welch | $t_{(13)}$ = 0.10, p = 0.92, t-test | $t_{(15)}$ = 0.40, p = 0.70, t-test |
| sIPSP tau | 4,91 ± 1,28 | 5,91 ± 1,57 | - | - | 5,96 ± 2,59 | 4,32 ± 1,29 | 4,95 ± 1,59 | 4,00 ± 0,96 | 6,85 ± 1,78 | 6,58 ± 2,06 | 4,15 ± 1,14 | 5,25 ± 2,36 | $t_{(15)}$ = 1.43, p = 0.17, t-test | - | U = 20, p = 0.23, M-W | $t_{(8)}$ = 1.06, p = 0.32, t-test | $t_{(13)}$ = 0.26, p = 0.80, t-test | $t_{(15)}$ = 1.20, p = 0.25, t-test |
| sEPSP fq | 0,53 ± 0,47 | 0,26 ± 0,17 | - | - | 0,08 ± 0,20 | 0,30 ± 0,28 | 0,39 ± 0,31 | 0,72 ± 0,41 | 0,11 ± 0,11 | 0,15 ± 0,21 | 0,23 ± 0,45 | 0,39 ± 0,58 | $t_{(8.716)}$ = 1.52, p = 0.16, Welch | - | U = 15, p = 0.08, M-W | $t_{(8)}$ = 1.44, p = 0.19, t-test | U = 20, p = 0.46, M-W | U = 20, p = 0.13, M-W |
| sEPSP amplitude | 17,07 ± 3,74 | 16,34 ± 3,32 | - | - | 15,58 ± 3,82 | 14,78 ± 2,66 | 13,30 ± 1,73 | 16,10 ± 2,68 | 13,66 ± 1,62 | 14,48 ± 1,61 | 13,96 ± 2,33 | 15,11 ± 2,90 | $t_{(15)}$ = 0.43, p = 0.68, t-test | - | $t_{(14)}$ = 0.49, p = 0.63, t-test | $t_{(8)}$ = 2.03, p = 0.08, t-test | $t_{(13)}$ = 0.97, p = 0.35 t-test | U = 16.50, p = 0.06, M-W |
| sEPSP 10-90 rise | 1,72 ± 0,23 | 1,68 ± 0,18 | - | - | 2,02 ± 0,26 | 1,85 ± 0,28 | 1,90 ± 0,29 | 1,67 ± 0,27 | 1,79 ± 0,25 | 1,69 ± 0,34 | 1,81 ± 0,15 | 1,76 ± 0,20 | U = 23, p = 0.24, M-W | - | $t_{(14)}$ = 1.27, p = 0.22, t-test | $t_{(8)}$ = 1.22, p = 0.26, t-test | U = 26, p = 0.95, M-W | $t_{(15)}$ = 0.63, p = 0.54 t-test |
| sEPSP half width | 4,12 ± 1,23 | 3,71 ± 0,58 | - | - | 4,05 ± 1,06 | 3,93 ± 1,14 | 3,53 ± 0,48 | 2,99 ± 0,46 | 4,42 ± 0,38 | 4,12 ± 0,82 | 3,81 ± 0,70 | 3,90 ± 1,16 | $t_{(15)}$ = 0.90, p = 0.38, t-test | - | U = 31, p = 0.94, M-W | $t_{(8)}$ = 1.79, p = 0.11, t-test | $t_{(13)}$ = 0.82, p = 0.43, t-test | $t_{(15)}$ = 0.19, p = 0.86, t-test |
| sEPSP tau | 2,88 ± 2,18 | 2,78 ± 1,18 | - | - | 3,18 ± 1,77 (7) | 3,92 ± 1,68 | 2,55 ± 0,87 | 2,06 ± 0,81 | 4,60 ± 1,81 | 2,59 ± 0,53 (8) | 3,10 ± 0,85 | 2,41 ± 1,15 (8) | $t_{(7.743)}$ = 0.99, p = 0.35, Welch | - | $t_{(13)}$ = 0.83, p = 0.42, t-test | $t_{(78)}$ = 0.89, p = 0.40, t-test | $t_{(5.652)}$ = 2.64, **p = 0.0410**, Welch | $t_{(14)}$ = 1.38, p = 0.19, t-test |

Continued:

| | A (8) | B (5) | C (7) | D (9) | E (4) | F (8) | G (6) | H (8) | I (7) | J (5) | K | L | | | | | | |
|---|---|---|---|---|---|---|---|---|---|---|---|---|---|---|---|---|---|---|
| mIPSC fq | 1,39 ± 2,66 | 1,05 ± 1,87 | 2,00 ± 1,22 | 2,32 ± 2,07 | 0,54 ± 1,02 | 1,66 ± 2,13 | 0,25 ± 0,39 | 0,39 ± 0,61 | 0,65 ± 0,31 | 0,97 ± 0,53 | - | - | U = 18, p = 0.83, M-W | $t_{(14)}$ = 0.36, p = 0.72, t-test | U = 2, **p = 0.0141**, M-W | U = 15, p = 0.28, M-W | $t_{(10)}$ = 1.31, p = 0.22, t-test | - |
| mIPSC amplitude | 14,88 ± 1,48 | 16,02 ± 3,81 | 15,56 ± 1,53 | 16,33 ± 4,12 | 14,34 ± 3,82 | 15,95 ± 1,60 | 12,72 ± 1,22 | 13,18 ± 1,66 | 15,39 ± 1,54 | 16,02 ± 1,97 | - | - | U = 18.5, p = 0.86, M-W | U = 31.5, p > 0.99, M-W | U = 8, p = 0.21, M-W | $t_{(12)}$ = 0.58, p = 0.58, t-test | $t_{(10)}$ = 0.63, p = 0.54, t-test | - |
| mIPSP 10-90 rise | 1,90 ± 0,20 | 1,77 ± 0,24 | 1,86 ± 0,09 | 1,71 ± 0,15 | 2,11 ± 0,43 | 1,79 ± 0,06 | 1,71 ± 0,25 | 1,83 ± 0,09 | 1,76 ± 0,07 | 1,81 ± 0,11 | - | - | $t_{(11)}$ = 1.08, p = 0.31, t-test | U = 8.5, **p = 0.0124**, M-W | $t_{(3.065)}$ = 1.44, p = 0.24, Welch | $t_{(6.075)}$ = 1.04, p = 0.34, Welch | $t_{(6.151)}$ = 0.91, p = 0.40, Welch | - |
| mIPSP half width | 6,19 ± 1,07 | 5,41 ± 0,92 | 6,14 ± 0,89 | 4,93 ± 0,74 | 7,13 ± 2,80 | 4,75 ± 2,06 | 5,65 ± 2,79 | 4,75 ± 2,06 | 6,17 ± 1,18 | 5,78 ± 0,97 | - | - | $t_{(11)}$ = 1.34, p = 0.21, t-test | $t_{(14)}$ = 2.97, **p = 0.0101**, t-test | $t_{(3.096)}$ = 1.08, p = 0.36, Welch | U = 18, p = 0.49, M-W | $t_{(10)}$ = 0.61, p = 0.56, t-test | - |
| mIPSP tau | 5,65 ± 1,15 | 5,60 ± 1,29 | 5,59 ± 1,44 | 4,57 ± 1,13 | 7,07 ± 2,62 | 5,25 ± 0,78 | 5,49 ± 2,49 | 4,81 ± 2,30 | 5,25 ± 1,41 | 5,05 ± 1,74 | - | - | $t_{(11)}$ = 0.08, p = 0.94, t-test | $t_{(14)}$ = 1.59, p = 0.13, t-test | $t_{(3.269)}$ = 1.36, p = 0.26, Welch | $t_{(12)}$ = 0.53, p = 0.60, t-test | $t_{(10)}$ = 0.22, p = 0.83, t-test | - |
| mEPSP fq | 0,10 ± 0,05 (7) | 0,34 ± 0,16 | 0,13 ± 0,07 | 0,20 ± 0,11 | 0,17 ± 0,13 | 0,18 ± 0,09 | 0,20 ± 0,15 | 0,30 ± 0,28 | 0,24 ± 0,09 | 0,29 ± 0,72 | - | - | $t_{(4.546)}$ = 3.17, **p = 0.0284**, Welch | $t_{(14)}$ = 1.48, p = 0.16, t-test | $t_{(10)}$ = 0.13, p = 0.90, t-test | $t_{(12)}$ = 0.76, p = 0.46, t-test | U = 14, p = 0.64, M-W | - |
| mEPSP amplitude | 11,44 ± 3,40 | 14,34 ± 3,28 | 13,22 ± 2,74 | 13,67 ± 2,27 | 15,76 ± 2,83 | 13,66 ± 1,37 | 12,79 ± 2,11 | 12,61 ± 1,66 | 14,43 ± 1,37 | 13,89 ± 2,61 | - | - | U = 11.5, p = 0.24, M-W | $t_{(14)}$ = 0.36, p = 0.72, t-test | $t_{(10)}$ = 0.24, p = 0.11, t-test | $t_{(12)}$ = 0.18, p = 0.86, t-test | $t_{(10)}$ = 0.47, p = 0.66, t-test | - |
| mEPSP 10-90 rise | 1,90 ± 0,20 | 1,79 ± 0,13 | 2,01 ± 0,28 | 1,65 ± 0,10 | 1,73 ± 0,93 | 1,67 ± 0,13 | 1,80 ± 0,27 | 1,79 ± 0,13 | 1,72 ± 0,10 | 1,65 ± 0,11 | - | - | $t_{(11)}$ = 1.14, p = 0.28, t-test | $t_{(7.118)}$ = 3.30, **p = 0.0128**, Welch | U = 10, p = 0.37, M-W | $t_{(12)}$ = 0.13, p = 0.90, t-test | $t_{(10)}$ = 1.23, p = 0.25, t-test | - |
| mEPSP half width | 4,43 ± 1,01 | 3,59 ± 0,90 | 4,27 ± 0,82 | 3,33 ± 0,45 | 4,33 ± 1,55 | 3,55 ± 0,51 | 3,45 ± 0,37 | 3,34 ± 0,40 | 3,95 ± 0,61 | 3,43 ± 0,60 | - | - | $t_{(11)}$ = 1.42, p = 0.18, t-test | $t_{(14)}$ = 2.94, **p = 0.0107**, t-test | $t_{(3.337)}$ = 0.98, p = 0.39, Welch | $t_{(12)}$ = 0.54, p = 0.60, t-test | $t_{(10)}$ = 1.46, p = 0.18, t-test | - |
| mEPSP tau | 4,77 ± 2,68 | 2,52 ± 1,10 | 2,62 ± 1,15 | 2,00 ± 0,64 (8) | 4,38 ± 3,30 | 3,05 ± 1,15 (7) | 2,58 ± 1,21 | 1,99 ± 0,39 | 2,82 ± 0,84 | 2,20 ± 0,50 | - | - | $t_{(10)}$ = 1.76, p = 0.11, t-test | **U = 6, p = 0.0093, M-W** | $t_{(3.491)}$ = 0.78, p = 0.49, Welch | $t_{(5.781)}$ = 1.15, p = 0.29, Welch | $t_{(10)}$ = 1.47, p = 0.17, t-test | - |

**Supplementary Table 10**. Sholl analysis - additional parameters

| Parameter | I RLN3 - mean ± SD (n) | | I pCCK- mean ± SD (n) | | II - mean ± SD (n) | | Two-way ANOVA (type III) - effects (p and F values with df) | | | Post hoc Tukey test (p and t values with df) | | | | | | | | |
|---|---|---|---|---|---|---|---|---|---|---|---|---|---|---|---|---|---|---|
| | | | | | | | | | | Ctrl | | | MS | | | Crtl vs. MS | | |
| | Ctrl (10) | MS (10) | Ctrl (12) | MS (12) | Ctrl (10) | MS (13) | Maternal separation | Type | Interaction | I RLN3 vs. I pCCK | I RLN3 vs. II | I pCCK vs. II | I RLN3 vs. I pCCK | I RLN3 vs. II | I pCCK vs. II | I RLN3 | I pCCK | II |
| Sum of intersections | 77.8 ± 26.9 | 60.3 ± 25.5 | 105.8 ± 28.5 | 71.0 ± 38.1 | 82.4 ± 29.0 | 75.6 ± 36.5 | $F_{(1,61)}$ = 6.44, **p = 0.01** | $F_{(2,61)}$ = 2.05, p = 0.14 | $F_{(2,61)}$ = 1.17, p = 0.32 | $t_{(61)}$ = 2.07, p = 0.10 | $t_{(61)}$ = -0.33, p = 0.94 | $t_{(61)}$ = 1.73, p = 0.20 | $t_{(61)}$ = 0.79, p = 0.71 | $t_{(61)}$ = -1.15, p = 0.49 | $t_{(61)}$ = -0.37, p = 0.93 | $t_{(61)}$ = 0.81, p = 0.42 | $t_{(61)}$ = 2.70, **p = 0.009** | $t_{(61)}$ = 0.51, p = 0.61 |
| Max. no. of intersections/ radius | 6.3 ± 1.7 | 4.9 ± 1.5 | 9.7 ± 2.4 | 6.3 ± 1.9 | 5.7 ± 2.1 | 6.2 ± 2.2 | $F_{(1,61)}$ = 8.56, **p = 0.005** | $F_{(2,61)}$ = 9.81, **p = 0.0002** | $F_{(2,61)}$ = 5.31, **p = 0.008** | $t_{(61)}$ = 3.96, **p = 0.0006** | $t_{(61)}$ = 0.68, p = 0.78 | $t_{(61)}$ = 4.67, **p = 0.0001** | $t_{(61)}$ = 1.69, p = 0.22 | $t_{(61)}$ = -1.50, p = 0.30 | $t_{(61)}$ = 0.23, p = 0.97 | $t_{(61)}$ = 1.58, p = 0.12 | $t_{(61)}$ = 4.12, **p = 0.0001** | $t_{(61)}$ = -0.54, p = 0.59 |

**Supplementary Table 11**. Sholl analysis

| Distance from soma - radius [µm] | I RLN3 - mean ± SD (n) | | I pCCK- mean ± SD (n) | | II - mean ± SD (n) | | Two-way ANOVA (type III) - effects (p and F values with df) | | | Post hoc Tukey test (p and t values with df) | | | | | | | | |
|---|---|---|---|---|---|---|---|---|---|---|---|---|---|---|---|---|---|---|
| | | | | | | | | | | Ctrl | | | MS | | | Crtl vs. MS | | |
| | Ctrl (10) | MS (10) | Ctrl (12) | MS (12) | Ctrl (10) | MS (13) | Maternal separation | Type | Interaction | I RLN3 vs. I pCCK | I RLN3 vs. II | I pCCK vs. II | I RLN3 vs. I pCCK | I RLN3 vs. II | I pCCK vs. II | I RLN3 | I pCCK | II |
| 10 | 3.1 ± 1.7 | 2.9 ± 1.1 | 3.6 ± 1.2 | 3.0 ± 1.8 | 2.7 ± 1.2 | 2.5 ± 1.1 | $F_{(1,61)}$ = 0.88, p = 0.35 | $F_{(2,61)}$ = 1.41, p = 0.25 | $F_{(2,61)}$ = 0.17, p = 0.85 | - | - | - | - | - | - | - | - | - |
| 20 | 4.1 ± 2.0 | 3.2 ± 0.6 | 5.8 ± 2.1 | 4.7 ± 1.4 | 3.5 ± 2.1 | 3.7 ± 1.3 | $F_{(1,61)}$ = 2.16, p = 0.15 | $F_{(2,61)}$ = 7.08, **p = 0.002** | $F_{(2,61)}$ = 0.98, p = 0.38 | $t_{(61)}$ = 2.33, p = 0.06 | $t_{(61)}$ = 0.81, p = 0.70 | $t_{(61)}$ = 3.18, **p = 0.007** | $t_{(61)}$ = 2.07, p = 0.11 | $t_{(61)}$ = -0.71, p = 0.76 | $t_{(61)}$ = 1.47, p = 0.31 | $t_{(61)}$ = 1.22, p = 0.23 | $t_{(61)}$ = 1.61, p = 0.11 | $t_{(61)}$ = 0.28, p = 0.78 |
| 30 | 4.4 ± 2.1 | 4.2 ± 1.8 | 6.2 ± 3.0 | 5.3 ± 2.3 | 3.4 ± 1.7 | 3.8 ± 1.6 | $F_{(1,61)}$ = 0.18, p = 0.67 | $F_{(2,61)}$ = 5.75, **p = 0.005** | $F_{(2,61)}$ = 0.59, p = 0.56 | $t_{(61)}$ = 1.93, p = 0.14 | $t_{(61)}$ = 1.04, p = 0.55 | $t_{(61)}$ = 3.02, **p = 0.01** | $t_{(61)}$ = 1.15, p = 0.49 | $t_{(61)}$ = 0.39, p = 0.92 | $t_{(61)}$ = 1.64, p = 0.24 | $t_{(61)}$ = 1.04, p = 0.30 | $t_{(61)}$ = -0.49, p = 0.62 | $t_{(61)}$ = 0.21, p = 0.84 |
| 40 | 4.8 ± 1.9 | 3.6 ± 0.7 | 6.1 ± 1.4 | 4.5 ± 1.4 | 3.7 ± 2.3 | 4.1 ± 1.8 | $F_{(1,61)}$ = 3.52, p = 0.07 | $F_{(2,61)}$ = 4.55, **p = 0.01** | $F_{(2,61)}$ = 1.63, p = 0.21 | $t_{(61)}$ = 1.82, p = 0.17 | $t_{(61)}$ = 1.50, p = 0.30 | $t_{(61)}$ = 3.39, **p = 0.004** | $t_{(61)}$ = 1.28, p = 0.41 | $t_{(61)}$ = -0.69, p = 0.77 | $t_{(61)}$ = 0.64, p = 0.80 | $t_{(61)}$ = 1.63, p = 0.11 | $t_{(61)}$ = 2.36, **p = 0.02** | $t_{(61)}$ = -0.55, p = 0.59 |
| 50 | 5.0 ± 1.4 | 3.6 ± 1.0 | 6.2 ± 1.9 | 4.8 ± 1.8 | 4.5 ± 2.5 | 4.4 ± 2.3 | $F_{(1,61)}$ = 4.39, **p = 0.04** | $F_{(2,61)}$ = 2.53, p = 0.09 | $F_{(2,61)}$ = 0.87, p = 0.43 | $t_{(61)}$ = 1.44, p = 0.33 | $t_{(61)}$ = 0.59, p = 0.83 | $t_{(61)}$ = 2.05, p = 0.11 | $t_{(61)}$ = 1.42, p = 0.34 | $t_{(61)}$ = -0.98, p = 0.59 | $t_{(61)}$ = 0.48, p = 0.88 | $t_{(61)}$ = 1.65, p = 0.10 | $t_{(61)}$ = 1.83, p = 0.07 | $t_{(61)}$ = 0.14, p = 0.89 |
| 60 | 4.9 ± 1.7 | 3.5 ± 0.8 | 7.0 ± 1.9 | 4.0 ± 1.6 | 3.8 ± 2.1 | 4.3 ± 2.1 | $F_{(1,61)}$ = 8.96, **p = 0.004** | $F_{(2,61)}$ = 4.72, **p = 0.01** | $F_{(2,61)}$ = 5.76, **p = 0.005** | $t_{(61)}$ = 2.78, **p = 0.02** | $t_{(61)}$ = 1.40, p = 0.35 | $t_{(61)}$ = 4.24, **p = 0.0002** | $t_{(61)}$ = 0.66, p = 0.79 | $t_{(61)}$ = -1.10, p = 0.52 | $t_{(61)}$ = -0.44, p = 0.91 | $t_{(61)}$ = 1.76, p = 0.08 | $t_{(61)}$ = 4.17, **p = 0.0001** | $t_{(61)}$ = -0.68, p = 0.50 |
| 70 | 4.1 ± 2.0 | 3.4 ± 1.1 | 6.8 ± 2.1 | 3.7 ± 1.7 | 4.3 ± 2.2 | 4.2 ± 2.0 | $F_{(1,61)}$ = 7.88, **p = 0.007** | $F_{(2,61)}$ = 3.43, **p = 0.04** | $F_{(2,61)}$ = 3.93, **p = 0.03** | $t_{(61)}$ = 3.26, **p = 0.005** | $t_{(61)}$ = -0.24, p = 0.97 | $t_{(61)}$ = 3.01, **p = 0.01** | $t_{(61)}$ = 0.33, p = 0.94 | $t_{(61)}$ = -0.94, p = 0.62 | $t_{(61)}$ = -0.64, p = 0.80 | $t_{(61)}$ = 0.82, p = 0.41 | $t_{(61)}$ = 3.98, **p = 0.0002** | $t_{(61)}$ = 0.18, p = 0.86 |
| 80 | 4.1 ± 2.5 | 3.6 ± 1.2 | 6.3 ± 2.3 | 3.4 ± 1.9 | 4.3 ± 2.3 | 4.2 ± 2.1 | $F_{(1,61)}$ = 5.30, **p = 0.03** | $F_{(2,61)}$ = 1.36, p = 0.27 | $F_{(2,61)}$ = 2.99, p = 0.06 | $t_{(61)}$ = 2.49, **p = 0.04** | $t_{(61)}$ = -0.21, p = 0.98 | $t_{(61)}$ = 2.26, p = 0.07 | $t_{(61)}$ = 0.20, p = 0.98 | $t_{(61)}$ = -0.63, p = 0.81 | $t_{(61)}$ = -0.88, p = 0.66 | $t_{(61)}$ = 0.53, p = 0.60 | $t_{(61)}$ = 3.40, **p = 0.001** | $t_{(61)}$ = 0.17, p = 0.87 |

Continued:

| | | | | | | | | | | | | | | | | | | |
|---|---|---|---|---|---|---|---|---|---|---|---|---|---|---|---|---|---|---|
| 90 | 3.4 ± 2.1 | 3.0 ± 1.2 | 6.0 ± 2.1 | 3.0 ± 1.8 | 3.8 ± 2.0 | 3.9 ± 2.1 | $F_{(1,61)}$ = 5.41, **p = 0.02** | $F_{(2,61)}$ = 2.53, p = 0.09 | $F_{(2,61)}$ = 4.45, **p = 0.02** | $t_{(61)}$ = 3.18, **p = 0.007** | $t_{(61)}$ = -0.47, p = 0.89 | $t_{(61)}$ = 2.69, **p = 0.03** | $t_{(61)}$ = 0.00, p = 1.00 | $t_{(61)}$ = -1.15, p = 0.49 | $t_{(61)}$ = -1.21, p = 0.45 | $t_{(61)}$ = 0.47, p = 0.64 | $t_{(61)}$ = 3.84, **p = 0.0003** | $t_{(61)}$ = -0.15, p = 0.89 |
| 100 | 3.3 ± 2.3 | 2.6 ± 1.3 | 5.0 ± 1.6 | 2.8 ± 1.7 | 3.8 ± 2.0 | 3.8 ± 1.6 | $F_{(1,61)}$ = 4.62, **p = 0.04** | $F_{(2,61)}$ = 1.89, p = 0.10 | $F_{(2,61)}$ = 4.45, p = 0.16 | $t_{(61)}$ = 2.23, p = 0.07 | $t_{(61)}$ = -0.63, p = 0.81 | $t_{(61)}$ = 1.57, p = 0.26 | $t_{(61)}$ = 0.31, p = 0.95 | $t_{(61)}$ = -1.66, p = 0.23 | $t_{(61)}$ = -1.42, p = 0.34 | $t_{(61)}$ = 0.88, p = 0.38 | $t_{(61)}$ = 2.98, **p = 0.004** | $t_{(61)}$ = -0.06, p = 0.95 |
| 110 | 3.2 ± 2.1 | 2.2 ± 1.4 | 4.8 ± 1.9 | 2.7 ± 1.4 | 3.7 ± 1.7 | 3.5 ± 1.5 | $F_{(1,61)}$ = 7.19, **p = 0.009** | $F_{(2,61)}$ = 2.44, p = 0.10 | $F_{(2,61)}$ = 2.09, p = 0.13 | $t_{(61)}$ = 2.27, p = 0.07 | $t_{(61)}$ = -0.66, p = 0.79 | $t_{(61)}$ = 1.57, p = 0.27 | $t_{(61)}$ = 0.65, p = 0.80 | $t_{(61)}$ = -1.89, p = 0.15 | $t_{(61)}$ = -1.29, p = 0.40 | $t_{(61)}$ = 1.33, p = 0.19 | $t_{(61)}$ = 3.15, **p = 0.003** | $t_{(61)}$ = 0.23, p = 0.82 |
| 120 | 3.1 ± 2.2 | 2.1 ± 1.4 | 4.1 ± 1.6 | 2.6 ± 1.4 | 3.0 ± 0.9 | 3.0 ± 1.5 | $F_{(1,61)}$ = 4.82, **p = 0.03** | $F_{(2,61)}$ = 1.23, p = 0.30 | $F_{(2,61)}$ = 1.41, p = 0.25 | $t_{(61)}$ = 1.49, p = 0.30 | $t_{(61)}$ = 0.15, p = 0.99 | $t_{(61)}$ = 1.57, p = 0.28 | $t_{(61)}$ = 0.73, p = 0.75 | $t_{(61)}$ = -1.39, p = 0.36 | $t_{(61)}$ = 0.67, p = 0.78 | $t_{(61)}$ = 1.45, p = 0.15 | $t_{(61)}$ = 2.38, **p = 0.02** | $t_{(61)}$ = 0.00, p = 1.00 |
| 130 | 2.8 ± 1.9 | 1.7 ± 1.3 | 3.5 ± 1.6 | 2.3 ± 1.6 | 3.1 ± 1.0 | 2.5 ± 1.6 | $F_{(1,61)}$ = 6.43, **p = 0.01** | $F_{(2,61)}$ = 0.20, p = 0.31 | $F_{(2,61)}$ = 0.27, p = 0.76 | $t_{(61)}$ = 1.08, p = 0.53 | $t_{(61)}$ = -0.44, p = 0.90 | $t_{(61)}$ = 0.62, p = 0.81 | $t_{(61)}$ = 0.98, p = 0.59 | $t_{(61)}$ = -1.32, p = 0.39 | $t_{(61)}$ = 0.34, p = 0.94 | $t_{(61)}$ = 1.63, p = 0.11 | $t_{(61)}$ = 1.89, p = 0.06 | $t_{(61)}$ = 0.88, p = 0.38 |
| 140 | 2.5 ± 1.4 | 1.6 ± 1.2 | 3.2 ± 1.5 | 2.3 ± 1.5 | 2.9 ± 1.0 | 2.1 ± 1.5 | $F_{(1,61)}$ = 6.82, **p = 0.01** | $F_{(2,61)}$ = 1.28, p = 0.28 | $F_{(2,61)}$ = 0.01, p = 0.99 | $t_{(61)}$ = 1.14, p = 0.50 | $t_{(61)}$ = -0.65, p = 0.79 | $t_{(61)}$ = 0.45, p = 0.89 | $t_{(61)}$ = 1.11, p = 0.51 | $t_{(61)}$ = -0.83, p = 0.69 | $t_{(61)}$ = 0.32, p = 0.95 | $t_{(61)}$ = 1.47, p = 0.15 | $t_{(61)}$ = 1.64, p = 0.11 | $t_{(61)}$ = 1.43, p = 0.16 |
| 150 | 2.0 ± 1.1 | 1.5 ± 1.1 | 2.9 ± 1.5 | 2.2 ± 1.5 | 2.6 ± 1.1 | 2.0 ± 1.4 | $F_{(1,61)}$ = 3.71, p = 0.06 | $F_{(2,61)}$ = 2.08, p = 0.13 | $F_{(2,61)}$ = 0.05, p = 0.95 | - | - | - | - | - | - | - | - | - |
| 160 | 1.7 ± 0.9 | 1.4 ± 1.0 | 3.0 ± 1.7 | 1.9 ± 1.6 | 2.4 ± 1.2 | 1.8 ± 1.4 | $F_{(1,61)}$ = 4.13, **p = 0.046** | $F_{(2,61)}$ = 2.50, p = 0.09 | $F_{(2,61)}$ = 0.47, p = 0.63 | $t_{(61)}$ = 2.26, p = 0.07 | $t_{(61)}$ = -1.17, p = 0.48 | $t_{(61)}$ = 1.04, p = 0.55 | $t_{(61)}$ = 0.90, p = 0.64 | $t_{(61)}$ = -0.65, p = 0.79 | $t_{(61)}$ = 0.27, p = 0.96 | $t_{(61)}$ = 0.50, p = 0.62 | $t_{(61)}$ = 1.98, p = 0.05 | $t_{(61)}$ = 1.12, p = 0.27 |
| 170 | 1.6 ± 0.8 | 1.4 ± 1.0 | 2.6 ± 2.2 | 1.8 ± 1.5 | 2.3 ± 1.3 | 2.0 ± 1.8 | $F_{(1,61)}$ = 1.19, p = 0.28 | $F_{(2,61)}$ = 1.35, p = 0.27 | $F_{(2,61)}$ = 0.20, p = 0.82 | - | - | - | - | - | - | - | - | - |
| 180 | 1.4 ± 1.0 | 1.5 ± 1.1 | 2.1 ± 1.7 | 1.6 ± 1.5 | 2.4 ± 1.5 | 1.8 ± 1.5 | $F_{(1,61)}$ = 0.83, p = 0.37 | $F_{(2,61)}$ = 1.20, p = 0.31 | $F_{(2,61)}$ = 0.34, p = 0.71 | - | - | - | - | - | - | - | - | - |
| 190 | 1.4 ± 1.0 | 1.1 ± 1.0 | 1.8 ± 1.9 | 1.4 ± 1.2 | 2.2 ± 1.4 | 1.8 ± 1.5 | $F_{(1,61)}$ = 1.28, p = 0.26 | $F_{(2,61)}$ = 1.51, p = 0.23 | $F_{(2,61)}$ = 0.01, p = 0.99 | - | - | - | - | - | - | - | - | - |
| 200 | 1.4 ± 1.0 | 1.1 ± 1.0 | 2.0 ± 2.4 | 1.3 ± 1.1 | 2.1 ± 1.4 | 1.8 ± 1.6 | $F_{(1,61)}$ = 1.32, p = 0.25 | $F_{(2,61)}$ = 1.17, p = 0.32 | $F_{(2,61)}$ = 0.18, p = 0.83 | - | - | - | - | - | - | - | - | - |
| 210 | 1.2 ± 1.0 | 1.1 ± 1.0 | 1.5 ± 1.4 | 1.1 ± 1.0 | 2.0 ± 1.4 | 1.7 ± 1.4 | $F_{(1,61)}$ = 0.84, p = 0.36 | $F_{(2,61)}$ = 1.99, p = 0.15 | $F_{(2,61)}$ = 0.09, p = 0.91 | - | - | - | - | - | - | - | - | - |
| 220 | 1.1 ± 1.0 | 1.1 ± 1.0 | 1.3 ± 1.3 | 0.8 ± 0.8 | 1.7 ± 1.1 | 1.5 ± 1.1 | $F_{(1,61)}$ = 1.11, p = 0.30 | $F_{(2,61)}$ = 1.76, p = 0.18 | $F_{(2,61)}$ = 0.43, p = 0.66 | - | - | - | - | - | - | - | - | - |
| 230 | 0.9 ± 1.0 | 1.0 ± 0.9 | 1.3 ± 1.3 | 0.7 ± 0.7 | 1.8 ± 1.3 | 1.1 ± 1.0 | $F_{(1,61)}$ = 2.80, p = 0.10 | $F_{(2,61)}$ = 1.46, p = 0.24 | $F_{(2,61)}$ = 1.01, p = 0.37 | - | - | - | - | - | - | - | - | - |
| 240 | 0.8 ± 0.9 | 0.9 ± 1.0 | 0.9 ± 0.5 | 0.6 ± 0.7 | 1.6 ± 0.8 | 1.2 ± 1.1 | $F_{(1,61)}$ = 1.17, p = 0.28 | $F_{(2,61)}$ = 3.54, **p = 0.04** | $F_{(2,61)}$ = 1.01, p = 0.38 | $t_{(61)}$ = 0.32, p = 0.95 | $t_{(61)}$ = -2.10, p = 0.10 | $t_{(61)}$ = -1.87, p = 0.16 | $t_{(61)}$ = -0.87, p = 0.66 | $t_{(61)}$ = -0.71, p = 0.76 | $t_{(61)}$ = -1.67, p = 0.23 | $t_{(61)}$ = -0.26, p = 0.79 | $t_{(61)}$ = 0.96, p = 0.34 | $t_{(61)}$ = 1.24, p = 0.22 |
| 250 | 0.7 ± 0.7 | 0.8 ± 0.9 | 0.8 ± 0.6 | 0.6 ± 0.7 | 1.6 ± 1.5 | 0.8 ± 0.9 | $F_{(1,61)}$ = 1.82, p = 0.18 | $F_{(2,61)}$ = 2.23, p = 0.12 | $F_{(2,61)}$ = 1.20, p = 0.31 | - | - | - | - | - | - | - | - | - |

Continued:

| | | | | | | | | | | | | | | | | | | |
|---|---|---|---|---|---|---|---|---|---|---|---|---|---|---|---|---|---|---|
| 260 | 0.7 ± 0.7 | 0.7 ± 0.7 | 0.8 ± 0.5 | 0.6 ± 0.7 | 1.3 ± 1.2 | 0.5 ± 0.7 | $F_{(1,61)}$ = 3.37, p = 0.07 | $F_{(2,61)}$ = 0.56, p = 0.57 | $F_{(2,61)}$ = 2.02, p = 0.14 | - | - | - | - | - | - | - | - | - |
| 270 | 0.7 ± 0.7 | 0.4 ± 0.5 | 0.8 ± 0.5 | 0.5 ± 0.7 | 1.2 ± 1.1 | 0.5 ± 0.7 | $F_{(1,61)}$ = 6.09, **p = 0.02** | $F_{(2,61)}$ = 0.92, p = 0.41 | $F_{(2,61)}$ = 0.82, p = 0.45 | $t_{(61)}$ = 0.17, p = 0.99 | $t_{(61)}$ = -1.58, p = 0.26 | $t_{(61)}$ = -1.48, p = 0.31 | $t_{(61)}$ = 0.33, p = 0.94 | $t_{(61)}$ = -0.21, p = 0.98 | $t_{(61)}$ = 0.14, p = 0.99 | $t_{(61)}$ = 0.95, p = 0.35 | $t_{(61)}$ = 0.87, p = 0.39 | $t_{(61)}$ = 2.48, **p = 0.01** |
| 280 | 0.7 ± 0.7 | 0.4 ± 0.5 | 0.8 ± 0.5 | 0.5 ± 0.7 | 0.9 ± 1.2 | 0.5 ± 0.7 | $F_{(1,61)}$ = 3.44, p = 0.07 | $F_{(2,61)}$ = 0.17, p = 0.84 | $F_{(2,61)}$ = 0.82, p = 0.47 | - | - | - | - | - | - | - | - | - |
| 290 | 0.6 ± 0.5 | 0.3 ± 0.5 | 0.8 ± 0.5 | 0.4 ± 0.5 | 0.8 ± 1.0 | 0.5 ± 0.7 | $F_{(1,61)}$ = 4.33, **p = 0.04** | $F_{(2,61)}$ = 0.46, p = 0.63 | $F_{(2,61)}$ = 0.01, p = 0.99 | $t_{(61)}$ = 0.55, p = 0.85 | $t_{(61)}$ = -0.71, p = 0.76 | $t_{(61)}$ = -0.18, p = 0.98 | $t_{(61)}$ = 0.43, p = 0.90 | $t_{(61)}$ = -0.61, p = 0.82 | $t_{(61)}$ = -0.18, p = 0.98 | $t_{(61)}$ = 1.06, p = 0.29 | $t_{(61)}$ = 1.29, p = 0.20 | $t_{(61)}$ = 1.27, p = 0.21 |
| 300 | 0.6 ± 0.5 | 0.3 ± 0.5 | 0.7 ± 0.5 | 0.6 ± 0.9 | 0.8 ± 1.0 | 0.4 ± 0.7 | $F_{(1,61)}$ = 2.33, p = 0.13 | $F_{(2,61)}$ = 0.36, p = 0.70 | $F_{(2,61)}$ = 0.33, p = 0.72 | - | - | - | - | - | - | - | - | - |
| 310 | 0.6 ± 0.5 | 0.3 ± 0.5 | 0.6 ± 0.5 | 0.4 ± 0.7 | 0.7 ± 0.9 | 0.4 ± 0.7 | $F_{(1,61)}$ = 2.69, p = 0.11 | $F_{(2,61)}$ = 0.11, p = 0.90 | $F_{(2,61)}$ = 0.09, p = 0.91 | - | - | - | - | - | - | - | - | - |
| 320 | 0.6 ± 0.5 | 0.3 ± 0.5 | 0.6 ± 0.7 | 0.4 ± 0.7 | 0.7 ± 0.9 | 0.5 ± 0.8 | $F_{(1,61)}$ = 1.88, p = 0.18 | $F_{(2,61)}$ = 0.19, p = 0.83 | $F_{(2,61)}$ = 0.05, p = 0.95 | - | - | - | - | - | - | - | - | - |

The symbol (-) indicates no post hoc analysis

**Supplementary Table 12**. Measured morphological parameters - descriptive statistics

| Morphological parameters | I RLN3 - mean ± SD (n) | | I pCCK- mean ± SD (n) | | II - mean ± SD (n) | |
|---|---|---|---|---|---|---|
| | Ctrl (10) | MS (10) | Ctrl (12) | MS (12) | Ctrl (10) | MS (13) |
| Nb_prim | 3.3 ± 1.2 | 3.0 ± 0.8 | 3.8 ± 1.0 | 3.9 ± 0.8 | 3.0 ± 0.8 | 3.0 ± 0.7 |
| Nb_bif | 5.5 ± 3.3 | 3.5 ± 2.0 | 8.2 ± 2.9 | 4.8 ± 1.7 | 6.7 ± 3.0 | 6.3 ± 5.3 |
| Nb_branch | 14.0 ± 6.4 | 9.6 ± 3.6 | 19.9 ± 5.6 | 13.2 ± 3.4 | 16.3 ± 6.5 | 15.2 ± 10.6 |
| Nb_tips | 8.5 ± 3.2 | 6.1 ± 1.7 | 11.8 ± 2.9 | 8.3 ± 1.8 | 9.6 ± 3.6 | 8.9 ± 5.4 |
| Max_branch | 2.7 ± 0.8 | 2.0 ± 1.1 | 3.7 ± 1.1 | 2.7 ± 1.2 | 3.3 ± 1.3 | 3.0 ± 1.6 |
| Tot_dendr [µm] | 970.7 ± 334.4 | 765.6 ± 313.3 | 1402.0 ± 422.5 | 912.0 ± 449.8 | 1081.0 ± 392.4 | 992.5 ± 488.1 |

**Abbreviations:** Max_branch, Maximal branch order; Nb_bif, Number of bifurcations; Nb_branch, Number of branches; Nb_prim, Number of primary dendrites; Nb_tips, Number of dendritic tips; Tot_dendr, Total dendritic length

**Supplementary Table 13.** Correlation matrix of the measured morphological parameters

| | **Nb_prim** | **Nb_bif** | **Nb_branch** | **Nb_tips** | **Max_branch** |
|---|---|---|---|---|---|
| **Nb_bif** | -0.01 | | | | |
| **Nb_branch** | 0.1 | 0.99* | | | |
| **Nb_tips** | 0.21 | 0.96* | 0.99* | | |
| **Max_branch** | -0.03 | 0.84* | 0.83* | 0.81* | |
| **Tot_dendr** | 0.07 | 0.70* | 0.71* | 0.70* | 0.62* |

* - Correlation significant at level p $\leq 0.05$

**Abbreviations:** Max_branch, Maximal branch order; Nb_bif, Number of bifurcations; Nb_branch, Number of branches; Nb_prim, Number of primary dendrites; Nb_tips, Number of dendritic tips; Tot_dendr, Total dendritic length

**Supplementary Table 14**. Correlation matrix of principal components and measured morphological parameters

| **Correlation matrix** | **PC1** | **PC2** |
|---|---|---|
| Nb_prim | 0.10 | -0,99* |
| Nb_bif | 0,98* | 0.10 |
| Nb_branch | 0,98* | -0.01 |
| Nb_tips | 0,97* | -0.12 |
| Max_branch | 0,89* | 0.14 |
| Tot_dendr | 0,80* | 0.00 |

**Abbreviations:** Max_branch, Maximal branch order; Nb_bif, Number of bifurcations; Nb_branch, Number of branches; Nb_prim, Number of primary dendrites; Nb_tips, Number of dendritic tips; Tot_dendr, Total dendritic length

**Supplementary Table 15.** Loadings of principal components

| Loadings | PC1 | PC2 |
|---|---|---|
| Nb_prim | 0.05 | -0.98 |
| Nb_bif | 0.47 | 0.10 |
| Nb_branch | 0.48 | -0.01 |
| Nb_tips | 0.47 | -0.12 |
| Max_branch | 0.43 | 0.14 |
| Tot_dendr | 0.38 | 0.002 |

**Abbreviations:** Max_branch, Maximal branch order; Nb_bif, Number of bifurcations; Nb_branch, Number of branches; Nb_prim, Number of primary dendrites; Nb_tips, Number of dendritic tips; Tot_dendr, Total dendritic length

**Supplementary Table 16.** Statistical analysis of principal components

| | Two-way ANOVA - effects (F and p values) | | | Post hoc Tukey test (t and p values) | | |
|---|---|---|---|---|---|---|
| | Maternal separation | Type | Interaction | C I RLN3 vs. MS I RLN3 | C I pCCK vs. MS I pCCK | C II vs. MS II |
| PC1 | $F_{(1, 61)} = 7.27$ **p = 0.009** | $F_{(2, 61)} = 3.64$ **p = 0.03** | $F_{(2, 61)} = 1.14$ p = 0.328 | $t_{(61)} = 1.52$ p = 0.13 | $t_{(61)} = 2.68$ **p = 0.009** | $t_{(61)} = 0.50$ p = 0.62 |
| PC2 | $F_{(1, 61)} = 0.0006$ p = 0.98 | $F_{(2, 61)} = 5.44$ **p = 0.007** | $F_{(2, 61)} = 0.40$ p = 0.67 | $t_{(61)} = -0.61$ p = 0.54 | $t_{(61)} = 0.67$ p = 0.51 | $t_{(61)} = 0.05$ p = 0.96 |
| | Post hoc Tukey test (p and t values) - continued | | | | | |
| | C I RLN3 vs. C I pCCK | C I RLN3 vs. C II | C I pCCK vs. C II | MS I RLN3 vs. MS I pCCK | MS I RLN3 vs. MS II | MS I pCCK vs. MS II |
| PC1 | $t_{(61)} = 2.32$ p = 0.06 | $t_{(61)} = -0.87$ p = 0.66 | $t_{(61)} = 1.41$ p = 0.34 | $t_{(61)} = 1.35$ p = 0.37 | $t_{(61)} = -2.04$ p = 0.11 | $t_{(61)} = -0.70$ p = 0.77 |
| PC2 | $t_{(61)} = -0.97$ p = 0.6 | $t_{(61)} = -0.87$ p = 0.66 | $t_{(61)} = -1.88$ p = 0.15 | $t_{(61)} = -2.24$ p = 0.07 | $t_{(61)} = -2.64$ **p = 0.03** | $t_{(61)} = -0.23$ p = 0.97 |

**Supplementary Table 17.** All counted cells

| All counted cells | | | | |
|---|---|---|---|---|
| NI part | Mean ± SD (n) | | p and t values with df | test |
| | Ctrl (5) | MS (6) | | |
| NI | 125 ± 20 | 126 ± 12 | $t_{(9)} = 0.07$, $p = 0.94$ | t test |
| NIc | 72 ± 13 | 75 ± 11 | $t_{(9)} = 0.51$, $p = 0.62$ | t test |
| NId | 54 ± 7 | 51 ± 9 | $t_{(9)} = 0.61$, $p = 0.56$ | t test |

**Supplementary Table 18.** Different mRNA species expression in the NI of MS and control rats

| mRNA species | Mean ± SD (% of counted cells/rat) (n) | | | | Two-way ANOVA - effects (p and F values with df) | | |
|---|---|---|---|---|---|---|---|
| | Ctrl (5) | | MS (6) | | Maternal separation | Localization | Interaction |
| | NIc | NId | NIc | NId | | | |
| vGAT1+ | 84 ± 5 | 41 ± 8 | 83 ± 11 | 49 ± 8 | $F_{(1,18)} = 1.004$, $p = 0.33$ | $F_{(1,18)} = 177.7$, $\mathbf{p < 0.0001}$ | $F_{(1,18)} = 1.76$, $p = 0.20$ |
| vGlut2+ | 19 ± 4 | 64 ± 10 | 19 ± 8 | 56 ± 9 | $F_{(1,18)} = 1.64$, $p = 0.22$ | $F_{(1,18)} = 142.1$, $\mathbf{p < 0.0001}$ | $F_{(1,18)} = 1.55$, $p = 0.23$ |
| RLN3+ | 47 ± 6 | 18 ± 4 | 46 ± 8 | 24 ± 4 | $F_{(1,18)} = 0.82$, $p = 0.38$ | $F_{(1,18)} = 101.5$, $\mathbf{p < 0.0001}$ | $F_{(1,18)} = 1.6$, $p = 0.22$ |
| CCK+ | 32 ± 8 | 3 ± 3 | 37 ± 8 | 3 ± 2 | $F_{(1,18)} = 1.25$, $p = 0.28$ | $F_{(1,18)} = 154.4$, $\mathbf{p < 0.0001}$ | $F_{(1,18)} = 0.67$, $p = 0.42$ |
| CRHR1+ | 72 ± 9 | 40 ± 11 | 81 ± 6 | 55 ± 9 | $F_{(1,18)} = 9.65$, $\mathbf{p = 0.006}$ | $F_{(1,18)} = 59.5$, $\mathbf{p < 0.0001}$ | $F_{(1,18)} = 0.41$, $p = 0.53$ |

| | | | | | | | |
|---|---|---|---|---|---|---|---|
| TrkA+ | 15 ± 3 | 18 ± 9 | 26 ± 9 | 29 ± 16 | $F_{(1,18)}$ = 5.31, **p = 0.03** | $F_{(1,18)}$ = 0.5, p = 0.49 | $F_{(1,18)}$ = 8E-07, p = 0.999 |

**Supplementary Table 19.** CRHR1 mRNA-expressing cells

| mRNA combination | Mean ± SD (% of counted cells/rat) (n) | | p and t values with df | test |
|---|---|---|---|---|
| | Ctrl (5) | MS (6) | | |
| **All CRHR1** | | | | |
| NI | 58 ± 8 | 71 ± 5 | $t_{(9)}$ = 3.14, **p = 0.01** | t-test |
| NIc | 69 ± 4 (4) | 83 ± 12 | U = 0, **p = 0.0095** | M-W |
| NId | 40 ± 11 | 54 ± 9 | $t_{(9)}$ = 2.31, **p = 0.047** | t-test |
| CRHR1 + vGAT1 | | | | |
| NI | 48 ± 7 | 57 ± 5 | $t_{(9)}$ = 2.2, p = 0.056 | t-test |
| NIc | 66 ± 10 | 73 ± 9 | $t_{(9)}$ = 1.16, p = 0.27 | t-test |
| NId | 24 ± 6 | 32 ± 4 | $t_{(9)}$ = 2.61, **p = 0.03** | t-test |
| CRHR1 + vGlut2 | | | | |
| NI | 13 ± 5 | 16 ± 9 | $t_{(9)}$ = 0.7, p = 0.5 | t-test |
| NIc | 8 ± 2 | 10 ± 8 | $t_{(5.88)}$ = 0.51, p = 0.63 | Welch |
| NId | 20 ± 10 | 26 ± 11 | $t_{(9)}$ = 0.93, p = 0.38 | t-test |
| CRHR1 + RLN3 | | | | |

| | | | | |
|---|---|---|---|---|
| NI | 33 ± 4 | 36 ± 6 | $t_{(9)} = 1.31$, $p = 0.29$ | t-test |
| NIc | 41 ± 1 (4) | 45 ± 8 | $t_{(5.29)} = 1.04$, $p = 0.34$ | Welch |
| NId | 18 ± 4 | 23 ± 4 | $t_{(9)} = 2.05$, $p = 0.07$ | t-test |
| CRHR1+ CCK | | | | |
| NI | 16 ± 7 | 21 ± 5 | $t_{(9)} = 1.61$, $p = 0.14$ | t-test |
| NIc | 26 ± 10 | 34 ± 9 | $t_{(9)} = 1.39$, $p = 0.2$ | t-test |
| NId | 2 ± 2 | 3 ± 1 | $t_{(9)} = 1.23$, $p = 0.25$ | t-test |

**Supplementary Table 20.** Mean area fraction of CRHR1 and TrkA-immunofluorescent dots per single cell

| **mRNA species** | **Mean ± SD (area fraction of counted cells/rat x10$^{-3}$)** | | | | **Two-way ANOVA - effects (p and F values with df)** | | |
|---|---|---|---|---|---|---|---|
| | **Ctrl (n)** | | **MS (n)** | | **Maternal separation** | **Localization** | **Interaction** |
| | **NIc (5)** | **NId (5)** | **NIc (6)** | **NId (5)** | | | |
| CRHR1+ | 2.46 ± 1.06 | 2.53 ± 1.06 | 2.97 ± 0.74 | 3.74 ± 0.63 | $F_{(1,17)} = 5.04$, **$p = 0.04$** | $F_{(1,17)} = 1.22$, $p = 0.29$ | $F_{(1,17)} = 1.22$, $p = 0.40$ |
| TrkA+ | 0.89 ± 0.34 | 2.02 ± 0.88 | 1.28 ± 0.71 | 3.76 ± 1.67 | $F_{(1,17)} = 5.89$, **$p = 0.03$** | $F_{(1,17)} = 16.80$, **$p = 0.0008$** | $F_{(1,17)} = 2.34$, $p = 0.14$ |

**Supplementary Table 21.** TrkA mRNA-expressing cells

| mRNA combination | Mean ± SD/ Median ± iqr (% of counted cells/rat) (n) | | p and t values with df | test |
|---|---|---|---|---|
| | Ctrl (5) | MS (6) | | |
| **All TrkA** | | | | |
| NI | 14 ± 2 (4) | 27 ± 12 | $t_{(5.35)}$ = 2.66, **p = 0.04** | Welch |
| NIc | 14 ± 1 (4) | 25 ± 13 | $t_{(5.03)}$ = 3.35, **p = 0.02** | Welch |
| NId | 18 ± 9 | 29 ± 16 | $t_{(9)}$ = 1.3, p = 0.23 | t-test |
| TrkA + vGAT | | | | |
| NI | 21 ± 3 (4) | 33 ± 22 | U = 0, **p = 0.0095** | M-W |
| NIc | 13 ± 1 (4) | 23 ± 7 | $t_{(5.35)}$ = 3.53, **p = 0.02** | Welch |
| NId | 9 ± 4 | 15 ± 9 | $t_{(9)}$ = 1.61, p = 0.14 | t-test |
| TrkA + vGlut2 | | | | |
| NI | 10 ± 2 (4) | 20 ± 13 | $t_{(5.54)}$ = 1.88, p = 0.11 | Welch |
| NIc | 1 ± 2 | 4 ± 3 | $t_{(9)}$ = 1.92, p = 0.09 | t-test |
| NId | 9 ± 2 (4) | 15 ± 10 | $t_{(5.69)}$ = 1.6, p = 0.16 | Welch |
| TrkA + RLN3 | | | | |
| NI | 6 ± 4 | 12 ± 4 | $t_{(9)}$ = 2.74, **p = 0.02** | t-test |
| NIc | 8 ± 4 | 14 ± 3 | $t_{(9)}$ = 2.77, **p = 0.02** | t-test |

| | | | | |
|---|---|---|---|---|
| NId | 4 ± 3 | 9 ± 6 | $t_{(9)}$ = 1.77, p = 0.11 | t-test |
| TrkA + CCK | | | | |
| NI | 3 ± 1 | 6 ± 2 | $t_{(9)}$ = 2.11, p = 0.06 | t-test |
| NIc | 4 ± 1 (4) | 9 ± 4 | $t_{(5.32)}$ = 2.91, **p = 0.03** | Welch |
| NId | 1 ± 1 | 1 ± 1 | $t_{(9)}$ = 0.53, p = 0.61 | t-test |
| TrkA + CRHR1 | | | | |
| NI | 10 ± 4 | 18 ± 7 | $t_{(9)}$ = 2.53, **p = 0.03** | t-test |
| NIc | 13 ± 4 | 22 ± 6 | $t_{(9)}$ = 2.83, **p = 0.02** | t-test |
| NId | 6 ± 3 | 13 ± 9 | $t_{(9)}$ = 1.69, p = 0.13 | t-test |

**Supplementary Table 22.** All RLN3 mRNA-expressing cells

| **mRNA Combination** | **Mean ± SD (% of counted cells/rat) (n)** | | **p and t values with df** | **test** |
|---|---|---|---|---|
| | **Ctrl (5)** | **MS (6)** | | |
| **All RLN3** | | | | |
| NI | 34 ± 4 | 37 ± 6 | $t_{(9)}$ = 0.86, p = 0.41 | t-test |
| NIc | 47 ± 6 | 46 ± 8 | $t_{(9)}$ = 0.20, p = 0.84 | t-test |
| NId | 18 ± 4 | 24 ± 4 | $t_{(9)}$ = 2.25, **p = 0.05** | t-test |
